# Colorectal cancer relies on an immunosuppressive cellular topography and genomic adaptations for establishing brain metastases

**DOI:** 10.1101/2025.11.14.688538

**Authors:** Anuja Sathe, Mengrui Zhang, Xiangqi Bai, Ji In Kang, Rithika Meka, Huiyun Sun, Susan M. Grimes, Aparajita Khan, Mingen Liu, Andrew S. Luksik, Michael Lim, Claudia K. Petritsch, Christopher M. Jackson, Hannes Vogel, Jeanne Shen, Melanie Gephart, Summer Han, Hanlee P. Ji

**Affiliations:** Division of Oncology, Department of Medicine, Stanford University School of Medicine, Stanford, CA, USA; Quantitative Sciences Unit, Department of Medicine, Stanford University School of Medicine, Stanford, CA, USA; Department of Neurosurgery, Stanford University School of Medicine, Stanford, CA, USA; Department of Neurosurgery, Johns Hopkins University School of Medicine, Baltimore, Maryland, USA; Stanford Cancer Institute, Stanford University School of Medicine, Stanford, CA, USA; Cancer Model Development Center, Stanford University School of Medicine, Palo Alto, USA; Department of Pathology, Stanford University, Stanford, CA, USA

## Abstract

Colorectal cancer (**CRC**) brain metastases have a poor prognosis and limited treatment options, including resistance to radiation therapy. Little is known about the molecular and cellular mechanisms that enable CRC tumor cells to adapt to the brain and establish a supportive tumor microenvironment. To address this gap we used spatial transcriptomics to analyze 51 CRC brain metastases. A subset had matched primary colon tumors and longitudinally paired metastatic resections before and after radiation treatment. We identified the critical spatial cellular features of the tumor epithelium and the surrounding tumor microenvironment that support metastatic growth in the brain. CRC brain metastases developed a stromal microenvironment with abundant fibroblasts and tumor-associated macrophages. A fibroblast–macrophage cellular neighborhood promoted angiogenesis, extracellular matrix remodeling, and immune suppression. Tumor cells showed local adaptations. In endothelial-rich regions, they were proliferative whereas in macrophage-rich regions, they were more differentiated and immune evasive. Compared with paired primary tumors, CRC brain metastases showed increased chromosomal instability, with activation of RNA-processing, stress response, and junctional remodeling pathways. After radiation treatment, resistant clones had increased epithelial–mesenchymal transition, while the immunosuppressive stroma remained intact. We identified tumor-derived *MIF*, *GDF15*, *PRSS3* and *SEMA3C* ligands and macrophage-derived *SPP1* that have the potential to affect multiple cell types in the metastatic niche. These ligand-receptor interactions drive angiogenesis, stromal activation and immune suppression. In a macrophage–tumor–fibroblast co-culture model, knockout of SPP1 in macrophages led to reduced expression of lipid-metabolism related genes and disrupted tumor-promoting interactions. Together, these results indicate that CRC growth in the brain is sustained by a specific cellular organization with immunosuppressive multicellular interactions.

## INTRODUCTION

Among the common epithelial cancers such as colorectal, lung and breast, metastasis is the leading cause of patient morbidity and mortality. One of the most consequential metastatic sites is the central nervous system (**CNS**). Patients with epithelial-derived brain metastases have dismal outcomes in terms of morbidity and mortality. Non-small cell lung cancers (**NSCLC**) and breast carcinomas frequently metastasize to the brain. While colorectal carcinomas (**CRCs**) can spread to the liver, lungs, and peritoneum (1), they also can spread to the brain with an estimated incidence of approximately 2% (2). Individuals with CRC brain metastasis (**CRC-BMets**) have survival ranging from 2 to 9.6 months (2, 3). Treating CRC-BMets is a challenge, even when combining surgery, radiation and chemotherapy – frequently they do not respond to radiation therapy (4, 5). Importantly, the incidence of patients with metastases is increasing because of the rising number of early onset CRC, defined as individuals younger than 50 (6).

Extrinsic and intrinsic cellular factors are required for a metastasis to establish and maintain itself in a new site. The extrinsic cellular factor comes from the foreign organ’s tumor microenvironment (**TME**). The brain represents a unique cellular milieu which is a challenge to seed and colonize compared to other organs (7). The blood brain barrier effectively isolates this CNS niche. The brain’s cellular microenvironment is composed of different neuronal and glial cell types. Importantly, this site is immunologically privileged, meaning that there are limitations on inflammation to prevent CNS tissue damage (8). Overall, cellular remodeling of the CNS niche is required for an epithelial cancer to grow in the brain.

The intrinsic cellular component comes from the properties of the tumor cells. Generally, very little is known about what genomic, molecular and cellular features enable the CRC tumor cells to establish themselves in the brain. Most metastatic CRCs have chromosomal instability (**CIN**), a molecular subtype with high levels of genomic instability and aneuploidy (9). In addition, CIN CRCs are microsatellite-stable (**MSS**) and as a result, not responsive to immune checkpoint blockade (10). While lung or breast cancer frequently metastasize to the brain, CRC has a lower propensity to establish tumors in the brain (11). Consequently, genomic and molecular analyses of brain epithelial metastases have focused on cancer types like lung or breast - both exhibit high rates of CNS involvement. Given its infrequent occurrence, CRC-BMets may have specific properties that distinguish them from the originating primary CRC and provide intrinsic adaptive advantages for growth in the brain.

Overall, there are very few studies of CRC-BMets. Prior genomic studies have primarily relied on conventional genome sequencing or single cell RNA-seq (**scRNA-seq**) to study the cellular composition of brain metastasis (12–15). While highly informative, this approach eliminates tissue structure. Samples are disaggregated into individual cells and as a result, there is a loss of the cellular spatial relationships in the native TME. There are no reports about this metastasis’ spatial cellular topography and related molecular features in this unusual niche.

Addressing this gap in our knowledge, we spatially characterized the CRC-BMet tumor cells and its TME. The spatial molecular and cellular features provided topographic, functional relationships of tumor cells to other proximal cell types and the adaptive genomic changes in the tumor and TME cells. We applied two complementary spatial transcriptomics assays to analyze a large set of CRC-BMets. One approach was a spatial in situ imaging assay that provided gene expression of specific markers at a single cell resolution. The other approach measured the spatial gene expression of nearly all genes. These spatial results allowed us to conduct an unbiased gene expression analysis, identify TME expression features and characterize their spatial relationships. Spatial information was particularly useful for an in-silico microdissection of specific cell types, like the tumor cells. This feature allowed us to examine genomic features exclusive to one cell type among the various TME components. We identified metastatic tumor cell adaptations including increased chromosomal instability and RNA processing. Notably, tumor cells adopted proliferative or quiescent states depending on their local microenvironment. *SPP1*-expressing macrophages were the dominant features of a fibroblast–macrophage niche. These macrophages are part of a critical cellular ‘hub’ that maintains an immunosuppressive microenvironment supporting tumor survival. We validated the importance of these *SPP1+* macrophages through an experimental cell culture system that replicated this cellular hub. Together, these results reveal a spatially organized multicellular program in which tumor and microenvironmental interactions sustain CRC-BMets.

## RESULTS

### Study overview

This sample set consisted of 51 CRC-BMets from 44 patients (**Supplementary Table 1-2**). Among this set, there were five matched primary CRC samples and one matched bladder metastasis. In addition, five different patients had longitudinal tissue samples from brain metastatic sites before and after radiation therapy. Lesions were distributed across the cerebellum, frontal, temporal, parietal and occipital lobes. Twenty-two tumors were confirmed through either mismatch repair protein or DNA molecular testing to be microsatellite stable (**MSS**), meaning that they lack a hypermutated phenotype related to loss of DNA mismatch repair (**MMR**) proteins. These MSS CRCs are not responsive to immune checkpoint agents. As we describe later, the remaining CRCs had additional genomic data indicating the molecular subtype. These patients had been treated with chemotherapy prior to surgical resection. We applied two complementary spatial transcriptomic approaches to these tumor samples (**Fig. 1A**). The first approach used imaging-based in situ transcriptomics with 480 genes (Xenium, 10X Genomics) on entire tissue sections (**Supplementary Table S3**). This method measures gene expression and spatial location at the resolution of single cells for each tissue section. There were 380 genes which were markers for delineating major immune (T, NK, B, plasma, macrophage) and stromal lineages (fibroblasts, endothelial). These genes also defined specific immunologic pathways (e.g., cytotoxicity, exhaustion, antigen presentation, etc.). We also included 100 genes that were markers of neurons, astrocytes, oligodendrocytes, microglia and colorectal cancer tumor cells. We used the in-situ approach on an overlapping subset of 11 CRC-BMets among the 51 samples, using adjacent tissue sections (**Supplementary Tables S2, S4**). This subset of 11 samples underwent analysis with both spatial assays. These 11 samples were selected based on tissue size to ensure complete lesion coverage within the capture area of each Xenium slide. These joint data sets on the same samples allowed us to make comparisons between the two spatial assays.

**Figure 1:**
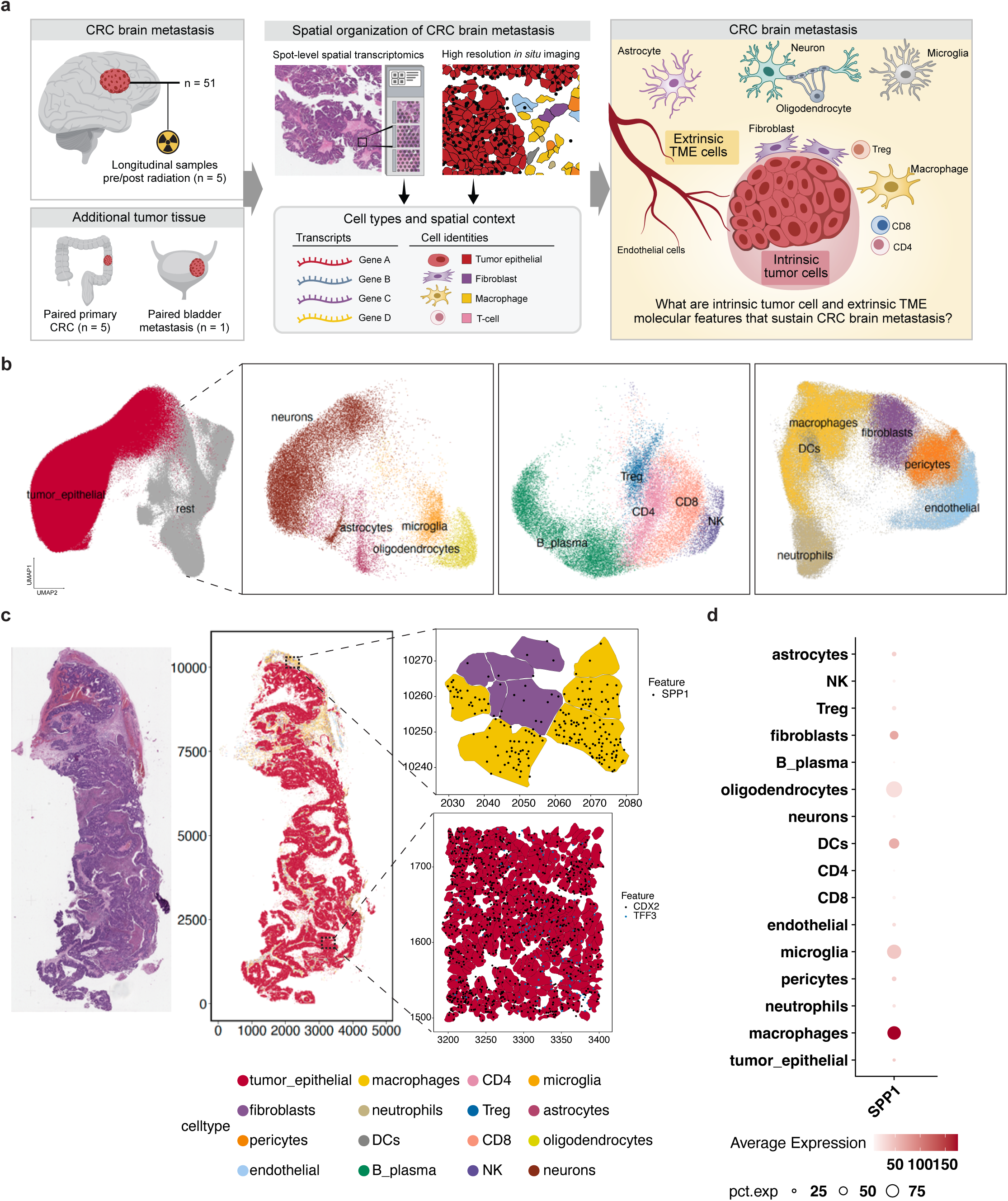
(a) Schematic representation of study design. (b) UMAP representation of Xenium data aggregated from all patients together with lineage specific sub clustering. (c) Example of paired H&E and spatially mapped cell types from P20, with zoomed in regions depicting respective cell types and transcripts. (d) Transcript counts of *SPP1* in respective cell lineages. Size of the dot is proportional to the percentage of cells expressing the gene.

The second approach used a sequencing-based spatial array (Visium, 10X Genomics) that topographically maps quantitative gene expression across the surface area of a tumor tissue section. The spatial array is composed of spots which provide gene expression data and its location coordinates across a tissue region. This approach measured the expression of 17,983 genes, representing a near complete transcriptome. With deconvolution of this expression data, specific cell types are identified with a fractional spatial assignment per given spot region. We quantitatively identified cell types and performed differential gene expression across different spatial regions. This approach was used to analyze all 51 CRC-BMets, the paired primary CRC and the paired bladder metastasis samples (**Supplementary Tables S2, S5**).

### Spatial in situ analysis identifies non-native immune and stromal cells in the brain TME

Prior studies have determined that the CRC TME, at its primary colon or rectal anatomic location, is a mixture of tumors cells, fibroblasts, macrophages, lymphocytes and others (16, 17). These TME cells provide a favorable niche through a combination of pro-tumor signaling, remodeling of the extracellular matrix (**ECM**) and suppressing anti-tumor immunity. These cell types, representing essential components of the colorectal TME, are scarce in normal brain parenchyma. Rather, the majority of immune and stromal cells in the brain are located in specialized tissue niches such as the meninges, a tissue covering the brain and the perivascular space that surrounds CNS blood vessels (18–20).

We analyzed 11 CRC-BMets with the spatial in-situ imaging method (Xenium), which provides gene expression at single cell resolution. These samples were used for this analysis based on their total regional area which maximized coverage for the in situ assay. The spatial imaging data was processed with results providing individual cell types with spatial assignment (**Methods**). We aggregated all cells for the 11 CRC-BMets samples, performed principal component analysis, conducted graph-based clustering and visualized the results with Uniform Manifold Approximation and Projection (**UMAP**) (**Methods**) (**Fig. 1b, Supplementary Fig. 1a**). Canonical marker genes were used to assign the cell type to each cluster (**Supplementary Fig. 1c-f**) (18, 19). We identified tumor, brain parenchymal (neurons, astrocytes, oligodendrocytes and microglia), stromal (fibroblasts, pericytes, endothelial cells), myeloid (macrophages, dendritic cells, neutrophils) and lymphocyte (T, NK, B, plasma) cell lineages from all samples (**Supplementary Fig. 1b**). For each cancer, we also obtained hematoxylin and eosin (**H&E**) stained images corresponding to the same tissue sections used for the spatial in situ assay (**Fig. 1c**). These histology images were evaluated by a pathologist. Using an image registration process, we aligned the H&E histopathology and the in-situ spatial cell assignments. Xenium cell type assignments matched the pathologist’s annotation (**Fig. 1c**).

The metastatic TME had desmoplastic fibroblasts, infiltrating macrophages and lymphocytes (20–22). Lymphocytes included naïve CD4 T cells, immunosuppressive T regulatory (**Treg**) cells, effector cytotoxic and exhausted CD8T cells, NK, B and plasma cells (**Supplementary Fig. 1d**). Fibroblasts expressed genes involved in ECM production (**Supplementary Fig. 1f**). Macrophages expressed tissue-remodeling and inflammatory genes that included *SPP1*, *MMP12*, *C1QC* and *CCL18* (**Fig. 1c, Supplementary Fig. 1c**). *SPP1,* which encodes osteopontin, is a secreted glycoprotein involved in matrix remodeling, cytokine signaling, and regulation of immune cell adhesion and migration (23). Its upregulation indicates activation of a macrophage state specialized for extracellular matrix remodeling and immunosuppressive signaling. Previous reports have shown that *SPP1*+ macrophages are indicators of tumor associated macrophages (**TAMs**) (24, 25). This result is in line with what prior studies described, namely that monocyte-derived TAMs were the dominant phagocytic population in brain metastases, not microglia (26, 27).

Macrophages showed the highest *SPP1* expression (Wilcoxon test, multiple-testing adjusted *p* ≤ 4.24e-29, log2FC = 0.96-5.35) among all cell types, including microglia (**Fig. 1d**). Notably, microglia can also express *SPP1* in neurogenerative diseases or following brain injury (28, 29). *SPP1*+ TAMs could either be derived from peripheral monocytes or resident microglia (30). As noted previously, several reports provide evidence pointing to these TAMs being monocyte derived (26, 27).

We also identified a distinct population of normal microglia, the brain’s tissue resident macrophages that are not monocyte derived. In comparison with macrophages, microglia had significantly higher expression of homeostatic microglial genes (*TMEM119*, *P2RY12*, *CX3CR1*) (Wilcoxon test, multiple-testing adjusted *p* <5e-8) (24) and lower expression of macrophage related genes (**Supplementary Fig. 1e**). As we describe later, these normal microglia were located in regions of the brain parenchyma distant to the tumor cells.

### Spatial transcriptomic assignments of cell types among the brain metastases

Next, we analyzed 51 CRC-BMets with the spatial sequencing-based transcriptomics assay (Visium) (**Supplementary Table S5**). As already noted, this approach provides a broader transcriptome-wide survey of gene expression levels over the area covered by the spatial array’s spot coordinates. To identify the cell composition, we used the program Cytospace on the spatial expression data (31). For a given spot region, this deconvolution method identifies the cell type and its fractional representation. For example, in a region that has exclusively CRC epithelium, this would be denoted as a 100% cell fraction. For determining cell fractions, this process requires a standardized single cell reference. Along these lines we constructed a scRNA-seq composite reference which included CRC-BMets, primary CRC and normal brain parenchyma (**Methods, Supplementary Fig. 2a-c**). Following deconvolution, we obtained the regional cell composition for all the CRCs and identified tumor epithelial, stromal (fibroblast, endothelial), immune (T/NK, B/plasma, macrophage) and brain parenchymal (neurons, astrocytes, oligodendrocytes, microglia) cells (**Fig. 2a, Supplementary Fig. 3a**). All the cell types were also identified from the in-situ Xenium analysis.

**Figure 2:**
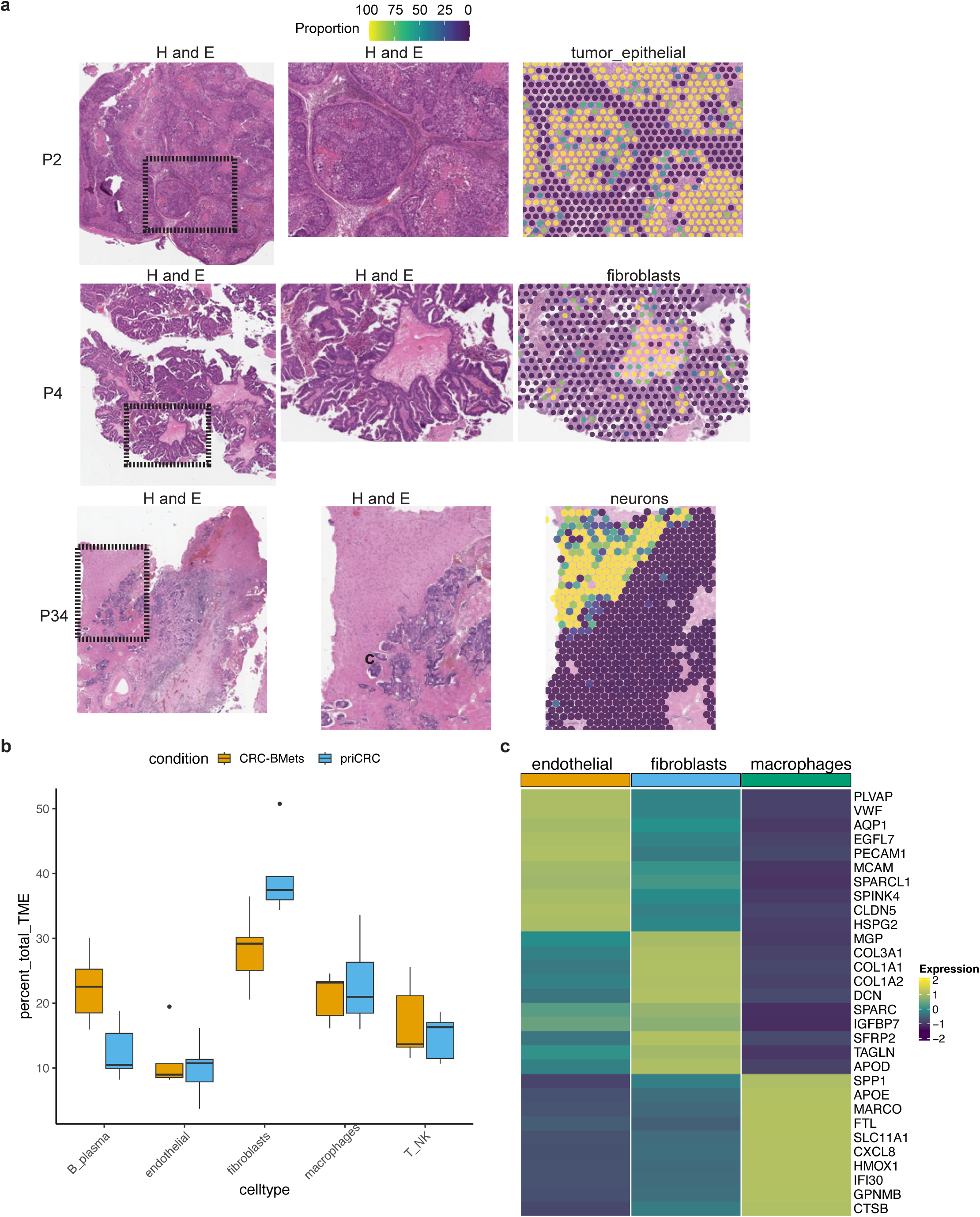
(a) Paired H&E and spot deconvolution proportions of Visium data for respective cell lineages in representative patients, together with zoomed in regions depicted in the inset. (b) Comparison of proportions between paired CRC-BMet and primary CRC samples from the same patient (Wilcoxon signed rank adjusted *p* = 0.14-0.58). (c) Scaled average expression of top differentially expressed genes in spots comprising respective cell lineages.

Five patients (P1, P4, P5, P40, P44) had matching paired primary CRC and CRC-BMets samples. The TME proportions of the shared macrophage, lymphoid and stromal cell types were similar and showed no statistical difference (Wilcoxon signed rank adjusted *p* = 0.14-0.58) (**Fig. 2b**). Overall, CRC-BMets contained immune and stromal cell types not native to the normal CNS parenchyma, with a fractional cell composition similar to what is observed among primary CRCs.

We validated the cell deconvolution results by comparing the average gene expression of cell types between the sequencing-based spatial gene expression (Visium) versus the spatial in situ imaging assay (Xenium). For this analysis, we focused on the 11 CRC-BMets with adjacent tissue sections and analyzed with both spatial assays (**Supplementary Table S2**). Between the two, there were 480 overlapping genes which were the basis for comparison. We used spot regions that had 100% fractional composition for each given cell type. This yielded cell type specific average gene expression measurements. For each sample, we compared the average gene expression obtained for the same cell type to the corresponding Xenium data. Overall, there was high concordance (average Spearman rho = 0.65, *p* < 7e-26) between the two spatial approaches results (**Supplementary Fig. 3b**).

### Differential gene expression among the cell types in the metastatic brain TME

For all 51 CRC-BMets, we evaluated the differential expression of the fibroblast, endothelial and macrophage populations. We leveraged the cell type assignments, using gene expression from regions with only a single cell type (i.e., 100% fraction). In essence, this is a in silico cell microdissection of specific cell types. These results were compared with expression profiles across all samples (Wilcoxon rank-sum test, log2FC > 0.25 and multiple-testing adjusted *p* < 0.05). Fibroblasts had upregulation of ECM related genes (**Fig. 2c, Supplementary Fig. 3c**). Macrophages had elevated expression of genes involved in matrix remodeling (*SPP1*, *CSTB*, *CTSH*, *LGMN*, *P4HA1, GPNMB*), lipid metabolism (*APOE*, *ABCG1*) and iron metabolism (*FTH1*, *FTL*) consistent with gene expression profiles of TAMs (23, 25). Spatial in situ analysis (Xenium) confirmed these results, also identifying desmoplastic fibroblasts and *SPP1*+ TAMs (**Fig. 1c-d, Supplementary Fig. 1c**). As we previously described, this subtype of TAM and cancer associated fibroblasts (**CAFs**) are a universal feature also among CRC liver metastasis (25).

### CRC epithelial adaptation to the metastatic brain niche

Comparing the primary colon site to the brain metastatic site, we evaluated the differences in CRC epithelium’s gene expression and identified upregulated pathways. The differential expression is a feature of how the tumor cells have adapted to the brain TME and are candidate factors contributing to metastatic tropism. We used the Visium data from five matching paired primary CRC and CRC-BMets (P1, P4, P5, P40, P44).

To ensure that we had gene expression data exclusively from CRC cells, we identified the regions that were composed entirely of cancer epithelium. This spatial gene expression data reflected a 100% tumor fraction. Then, we compared gene expression between the primary colon and the matched brain metastases tumor cells (**Fig. 3a, Supplementary Table 6**). Differential expression was performed using Model-based Analysis of Single-cell Transcriptomics (**MAST**) (32). This method incorporates variation from both biological and technical sources, modelling gene expression as a mixed-effects regression model. For this analysis, the number of detected genes per spatial spot region was included as a covariate to control for sequencing depth, and patient was modelled as a random effect to account for inter-patient variability. This design allowed the identification of genes whose expression differences were consistently observed across all five CRC pairs (**Supplementary Fig. 4a**). The criteria for differential expression included a log2FC ≥ 0.5 and multiple-testing adjusted *p* ≤ 0.05.

**Figure 3:**
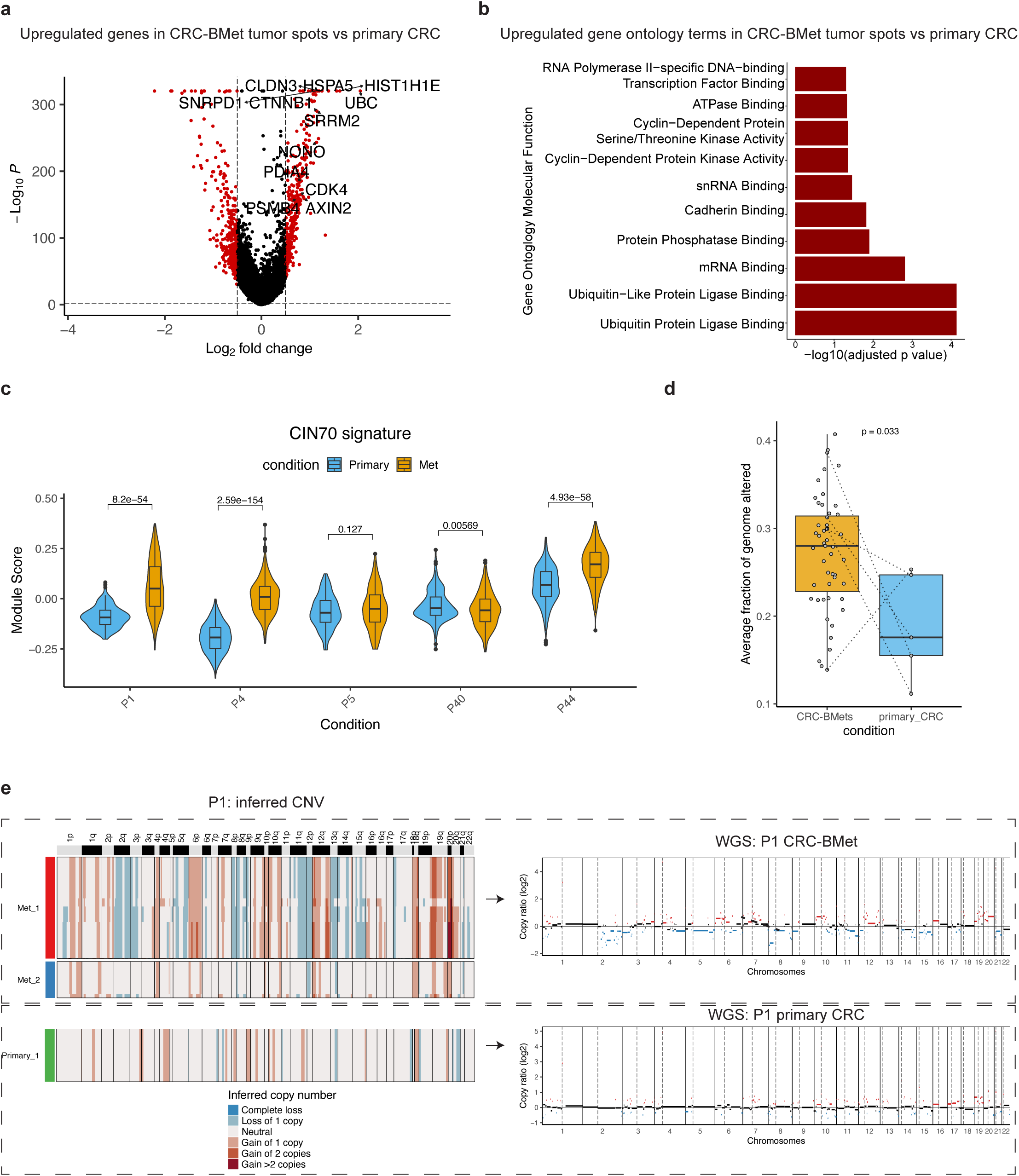
(a) Volcano plot highlighting upregulated genes in pure tumor spots from CRC-Bmets compared to paired primary CRC. (b) Gene ontology analysis on upregulated genes in CRC-BMet tumor spots (c) Comparison of CIN70 signature in pure tumor spots from CRC-Bmets compared to paired primary CRC from respective samples with t-test *p*. (d) Average inferred fraction of genome altered in pure tumor spots from CRC-Bmets compared to paired primary CRC with t-test *p*. Dotted lines connect paired samples from the same patient. (e) Inferred CNV profiles from pure tumor spots from paired CRC-Bmets and primary CRC from P1, with suffix indicating subclone, together with CNV profiles from WGS.

Compared to the colon primary tumors, CRC-BMet tumor cells significantly upregulated the expression of RNA spliceosome components and nuclear body–associated factors. These genes included several small nuclear ribonucleoproteins (*SNRNP70, SNRPD1, SNRPC*) and splicing regulators (*SRRM2, SRSF3, SRSF7, TRA2B*). There were also differentially expressed genes linked to nuclear speckles and paraspeckles (*NONO, HNRNPC, HNRNPR, PRRC2A*). These nuclear bodies are composed of interacting RNA-binding proteins (**RBPs**) and long noncoding RNAs (33). Nuclear speckles and paraspeckle regulate gene expression through multiple mechanisms. Spliceosome and speckle pathways have been implicated in sustaining cell type plasticity and stress tolerance in cancer (34, 35).

Epithelial junction remodeling was upregulated in CRC-BMets. This biological process involves assembly of the junctional complexes that mediate adhesion with neighboring cells and the ECM (36). In the brain, tumor cells had significantly increased expression of multiple genes from the claudin family (*CLDN3*, *CLDN4*, *CLDN7*). Using immunohistochemistry (**IHC**) we confirmed the upregulation of CLDN3 protein in CRC-BMets compared to primary CRC (**Supplementary Fig. 4b**). Claudin overexpression results in disruption of epithelial tight junctions and contributes to metastasis (37). We also observed overexpression of *AXIN2* and *CTNNB1* related to Wnt signaling in the CRC-BMets compared to primary CRC.

Protein metabolism and proteostasis were altered among the CRC epithelial cells in the brain. Proteostasis represents the general homeostatic state of protein synthesis, folding and degradation. The CRC-BMet tumor cells significantly upregulated the proteasome subunits (*PSMA1, PSMB4, PSMB7*), chaperones (*HSPA5, DNAJB1, VCP*) and unfolded protein response (**UPR**) regulators (*XBP1, PDIA4, PDIA6*). Together these findings indicate activation of proteostasis and endoplasmic reticulum (**ER**) stress pathways. The dysregulation of these processes supports tumor cell survival in the metabolic conditions of the brain niche (38). Enrichment of these pathways in CRC-BMet tumor cells compared to the primary CRC was confirmed by gene ontology analysis (39) (**Fig. 3b**). These gene expression features were consistently present among all the brain metastases (**Supplementary Fig. 4a**).

We had additional evidence that these upregulated pathways were specific to brain metastases. We had gene expression results from a paired bladder metastasis and a CRC-BMet, both from the same patient (P2). We identified differentially expressed genes among the tumor cells from the two sites (Wilcoxon test, log2FC = 0.5, multiple-testing adjusted *p* ≤ 0.05). Compared to the bladder site, the brain metastatic tumor cells showed upregulation of spliceosome and RNA-processing factors (*EIF4B, SNRPC, PRPF8, SRSF3*), WNT modulators (*CTNNB1, NOTUM, NKD1, SP5*), stress-response (*DDIT4, HSP90AB1*) and proliferation (*CCND1, CDK4*) (**Supplementary Fig. 4c**). These results were consistent with the CRC-BMet tumor cells having increased expression of RNA processing, WNT signaling, and junctional remodeling compared to primary CRC. These molecular features represent adaptations specific to the brain niche rather than generic features of CRC metastasis.

Next, we determined if these pathways were also upregulated among the other CRC-BMets. For this analysis, we determined the Hallmark gene set pathway activity across all CRC-BMets (40). Clustering revealed two patterns of pathway activation (**Supplementary Fig. 5a**). The first was defined by the cell cycle, MYC activity, ongoing DNA repair, the unfolded protein response and WNT signaling, consistent with proliferative and stress-adapted states. The second showed increased apical junction, TGF-β signaling and inflammatory pathways which indicate epithelial remodeling and cytokine-driven interactions. Together, these results show that colorectal cancer cells take on adaptive features to the brain niche including: RNA processing, WNT signaling, junctional remodeling, proteostasis and ER stress tolerance.

### Chromosomal alterations and aneuploidy in CRC brain metastases

We further examined the genomic properties of these metastatic tumor cells from their spatial assignments. We determined if the CRC-BMets had chromosomal arm imbalances, which are the hallmarks of aneuploidy. To infer chromosomal level copy number variations (**CNV**) we extracted the gene expression data from spatial regions that were composed entirely of cancer epithelium (100% tumor fraction). Then we used the inferCNV algorithm (41, 42) (**Methods**), which estimates CNVs based on average gene expression of large chromosome segments. Previously, we demonstrated that this approach accurately identifies chromosomal imbalances from single cell and spatial genomic data (43, 44). In addition, we calculated the inferred fraction of genome altered (**FGA**), which is a measurement of the proportion of DNA segments with copy number gains or losses (45). Finally, we determined the enrichment of the CIN70 signature for each tumor rich region (46). This signature correlates with the degree of aneuploidy present in a given cancer. Together, the inferred FGA value and CIN70 signature score provided additional quantitative measures of genomic instability.

We determined the chromosome and genomic instability properties among the five matched metastatic and the primary CRCs (P1, P4, P5, P40, P44). Compared to the paired primary CRCs, brain metastatic tumors had significantly higher FGA (t-test *p*=0.033) and CIN70 values (t-test *p* range 0.127-2.6e-154), indicating higher degrees of genomic instability (**Fig 3c-d**). The CRC-BMet tumor cells had more chromosomal arm imbalances compared to the primary CRCs. In three or more brain metastases, there were more amplifications in 6p, 7p, and 20q, and deletions in 15q compared to the paired primary CRCs (**Fig. 3e, Supplementary Fig. 6a-d**). These CNV patterns were distributed in distinct subclones suggestive of clonal diversity in these lesions. We conducted whole-genome sequencing (**WGS**) from all paired CRC-BMet and primary CRC samples (**Fig. 3e, Supplementary Fig. 2a-d**). The WGS results confirmed the presence of the arm imbalances which were identified from the spatial data.

We compared the paired CRC-BMet and bladder metastasis from P2. The CRC-BMet tumor cells had significantly higher CIN70 signature (t-test *p*=5.7e-11) and inferred FGA (t-test *p*=2.89e-58) compared to the paired bladder metastasis (**Supplementary Fig. 7a-b**). CRC-BMet tumor cells shared several CNV events with the bladder metastasis consistent with a shared origin (**Supplementary Fig. 7d**). However, the CRC-BMet also had additional chromosomal imbalances including gains in 10p, 12p, 12q, 15q, 17q, 2q and deletions in 11p, 13q, 19q, 20p, 21q, 3q, 4q, 9p and 9q. The increased number of arm imbalances was a notable feature for the brain metastasis.

Next, we determined the chromosomal arm imbalances for all CRC-BMets, excluding the primary CRCs and the bladder metastasis. Seventy percent of the CRC-BMets had recurrent events which included gains (2p, 7p, 6p, 19p, 19q, 8q) and losses (8p and 18p) (**Supplementary Fig. 7d**). These results are consistent with a recent study examining specific CNV alterations which occur frequently in CRC-BMets (9). Among the CRC-BMets, inferred focal copy number changes included amplifications of *MYC* (57%), *CCND1* (51%) and the S100 gene family (*∼*70%) (**Supplementary Table 7**). These genes are established drivers of CRC metastasis (9). Collectively, these results show that CRC-BMets have elevated chromosomal imbalances distinct from the primary CRC and are indicators of ongoing metastatic clonal evolution. Also, this result indicates that the tumors are of the CIN molecular subtype which are associated with being microsatellite stable (9).

### Cell neighborhoods of CRC brain metastases

As previously described, the CRC-BMet TME contains stromal and immune cell types not present in the normal brain parenchyma (**Fig. 1**, **2**). To understand how these cells spatially localize in relationship to each other, we determined the cell neighborhood (**CN**) properties from the Xenium data (47). CNs are quantitative indicators of the local spatial cellular contents and its organization; they provide specific information about interactions among the tumor, stromal and immune cells. We used the imcRtools software (48), which has been extensively used in studies implementing CN analysis (49–51). We quantified the proportion of all cell types within a fixed radius of 65 μm surrounding each individual cell. This radius was selected since it corresponds to the size of each Visium spot, enabling a later comparison of our findings from Xenium. Then, we used k-means clustering to group neighborhoods with a similar composition. Each cluster defined the CN and the interacting cell types in the metastatic TME (**Fig. 4a**).

**Figure 4:**
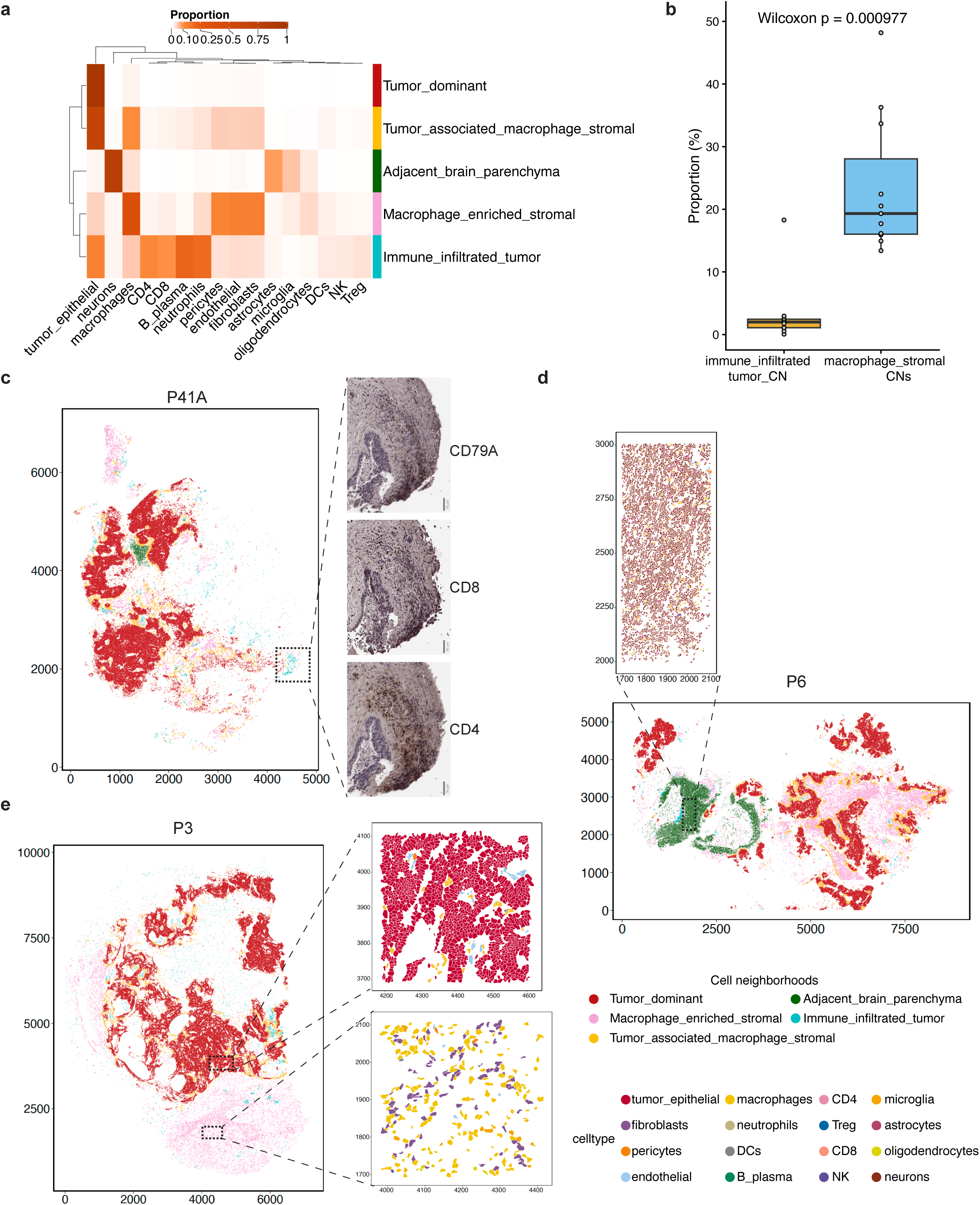
(a) Hierarchical clustering of cellular proportions across cell neighborhoods. (b) Proportions of immune infiltrated CN or macrophage-stromal CNs (sum of macrophage enriched stromal and tumor associated macrophage stromal CNs) across samples. (c-e) Spatial representation of cellular neighborhoods in respective samples. Zoomed inset in b depicts IHC for respective proteins performed in adjacent sections, insets in d and e depicts cell types.

There were five categories of CNs. Each one was labelled based on the proportions of their constituent cells in the TME. We identified i) tumor dominant, ii) immune-infiltrated tumor, iii) macrophage enriched stromal, iv) tumor-associated macrophage stromal and v) adjacent brain parenchyma CNs (**Fig. 4a, Supplementary Fig. 8a**). The tumor dominant CN was composed of tumor epithelial cells, making up 96% of all cells. The macrophage enriched stromal CN was dominated by macrophages (38%) together with endothelial (14%), fibroblast (14%) and pericyte (13%) stromal components. The tumor associated macrophage stromal CN also contained macrophages and stromal cells but in addition contained tumor epithelial cells (66%). The immune infiltrated tumor CN contained tumor epithelial cells (13%) together with CD4 (11%), CD8 (10%) and B/plasma (21%) cells. To verify the immune infiltrated CN with a different method, we used IHC to determine the protein expression of a set of immune cell markers (CD4, CD8 and CD79A) on adjacent sections of CRC-BMet that had been analyzed with Xenium (**Fig. 4c, Supplementary Fig. 4b**). The IHC results confirmed this CN. Finally, the adjacent brain parenchymal CN contained the highest proportion of neurons (78%) together with astrocytes (8%). This CN contained almost no fibroblasts (0.21%), indicating that the fibroblast-rich stroma is associated with cancer cells rather than being a property of the surrounding brain. Microglia were most abundant in this CN (5.6%) compared to all other CNs (0.01-1.8%), reinforcing that normal homeostatic microglia are largely confined to the adjacent brain parenchyma rather than tumor-associated regions.

We evaluated the spatial relationship among the different CNs by visualizing their distribution across all samples (**Fig. 4c-d, Supplementary Fig. 6b**). The tumor dominant CNs were bordered by tumor associated macrophage-stromal CNs, with intervening macrophage enriched stromal CNs. The immune-infiltrated tumor CN, in which lymphocytes and macrophage cells are intermingled with tumor cells, were rare (0.1–2.9% of total tissue), with the exception of one CRC-BMet (P44) as an outlier (**Supplementary Fig. 8a**). The overall proportions of immune-infiltrated CN were significantly lower (Wilcoxon test, *p*=9e-4) than CNs with macrophages and stromal cells (macrophage enriched stromal and tumor-associated macrophage stromal) (**Fig. 4b**). This result indicated that dense intratumoral lymphocyte neighborhoods are uncommon, consistent with CRC-BMets and their TME excluding lymphocytes. Overall, these results revealed coordinated spatial organization among tumor cells, macrophages and stromal cells in CRC-BMets.

### Macrophages, fibroblasts and tumor cells are spatially associated in the brain metastatic niche

Using the spatial gene expression data (Visium) we extended our analysis of CN spatial organization to all CRC-BMets. We used MistyR, a random-forest method that quantifies spatial relationships from spatial gene expression data (52). MistyR models how the abundance of one cell type predicts the abundance of another cell type – this model provides an ‘importance’ score that quantifies the strength of these relationships. For each sample, we used the cell type fractions for any given spatial spot radius of 65 μm. The MistyR analysis showed that the abundance of tumor, fibroblast and macrophage cells positively predicted one another in more than 70% of samples, whereas neuronal and other CNS cells predicted each other but not fibroblasts (**Supplementary Fig. 9a–b**). The analysis showed the same associations even with expanded spatial spot radius of 250 μm (**Supplementary Fig. 9c**). Collectively, these results indicated that CRC-BMets are organized epithelial tumor regions embedded within macrophage- and fibroblast-rich stroma.

### Gene expression features of the spatial domains among CRC-BMets

Gene expression programs involve sets of genes activated together to support specific biological pathways. We determined whether specific gene programs correlated with the spatial organization of the different TME cells. This analysis was based on the spatial gene expression data (Visium) from all CRC-BMet samples. We used the Banksy algorithm, which performs spatially aware clustering by integrating local spatial proximity and gene expression similarity (53). The resulting clusters represent a contiguous region of cells with distinct gene expression programs - a given cluster is referred to as a ‘spatial domain’. Unlike cell neighborhood analysis, which identifies only the colocalization of cell types, spatial domains define tissue regions by integrating transcriptional similarity with spatial proximity. Compared to methods that rely solely on transcription-based clustering, this approach captures transcriptional heterogeneity within spatially proximal regions. Hence, this approach enables the assessment of how spatial localization influences cell gene expression phenotypes and function (54).

We identified eight spatial domains that were present in nearly all CRC tumors (**Fig. 5a-b, Supplementary Fig. 10a-b**). Each domain was defined based on its cellular composition and the expression of tumor and TME hallmarks of cancer biological pathways (**Fig. 5a-c**). This included pathways related to tumor progression and microenvironmental remodeling (55) (**Fig. 5c**). The eight domains were present from 70.6 to 100% across all of the brain metastases (**Supplementary Fig. 10a**).

**Figure 5:**
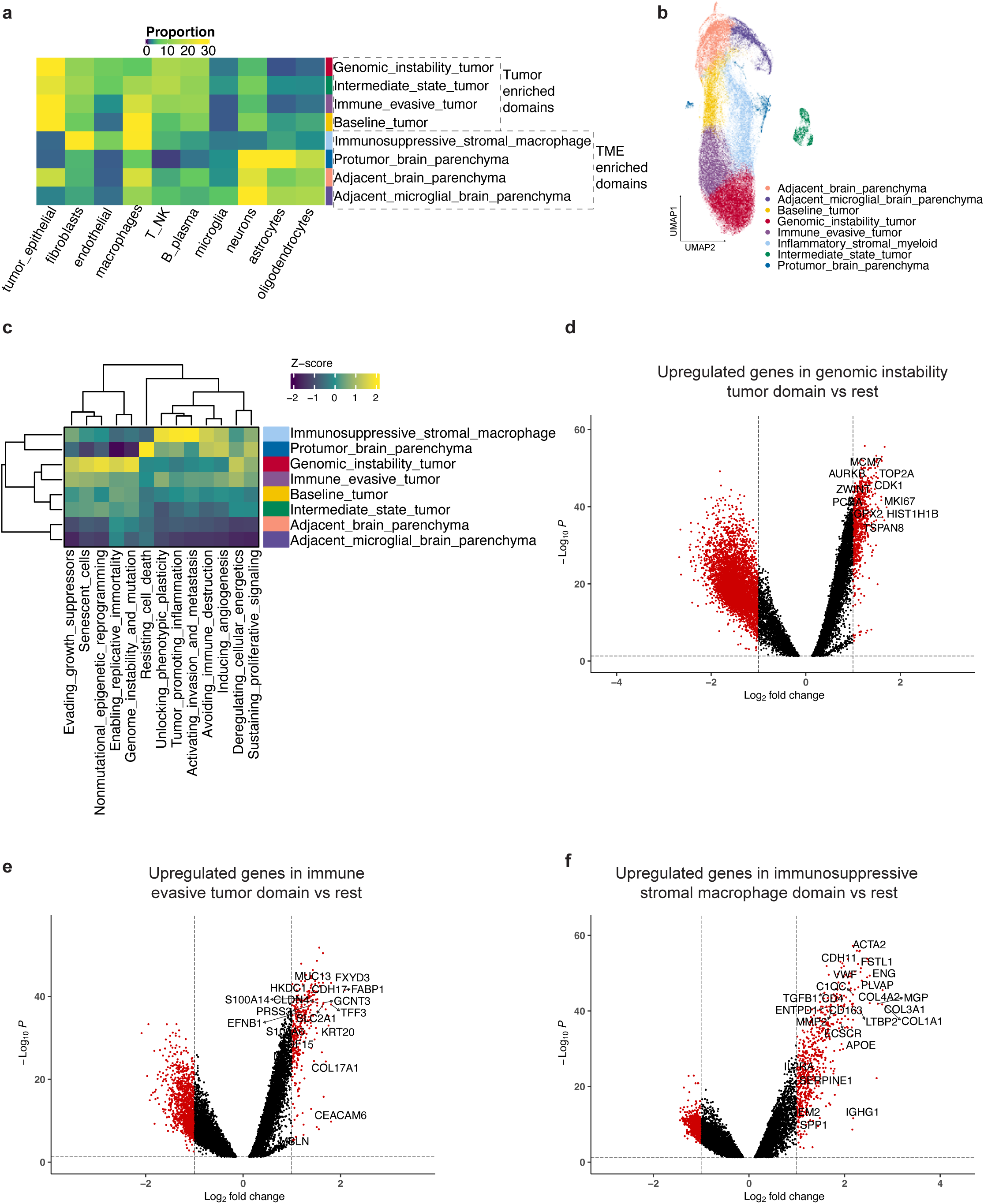
(a) Cellular composition of spatial domains. (b) UMAP representation of spatial domains. (c) Scaled average pathway activity across spatial domains. (d-f) Volcano plots highlighting upregulated genes in (d) genomic instability tumor domain or (e) immune evasive tumor domain or (f) immunosuppressive stromal macrophage domain compared to other domains.

The first four spatial domains were enriched for tumor epithelial cells with varying proportions of other TME cell types (**Fig. 5a**). These domains were labelled based on the most prominent hallmark gene expression program (**Fig. 5c**). They included: i) baseline tumor, ii) genomic instability, iii) immune evasive and iv) intermediate state. This result indicated that tumor cells adopt spatially restricted expression states shaped by the surrounding microenvironment. A fifth spatial domain was defined by v) immunosuppressive fibroblasts and macrophages (**Fig. 5a**). This domain showed coordinated upregulation of immune suppression pathways including tumor promoting inflammation, activating invasion and metastasis, avoiding immune destruction and inducing angiogenesis (**Fig. 5c**).

The sixth, seventh and eighth spatial domains had brain parenchymal cell types including neurons, astrocytes and oligodendrocytes (**Fig. 5a**). The sixth brain parenchymal domain had vi) pro-tumor properties and showed selective activation of angiogenic, inflammatory, and survival programs that can promote tumor growth (**Fig. 5c**). The seventh and eighth brain parenchymal domains lacked pro-tumor features and were indicative of physiologic brain parenchyma. Together, these spatial domains defined spatially distinct gene expression programs that are indicators of tumor adaptation, immunosuppression and tissue remodeling in the metastatic brain environment.

### Gene expression features of the brain metastatic spatial domains

We examined the specific gene expression features underlying each spatial domain. For each CRC-BMet and a given domain, the gene expression was aggregated across the spatial regions - this procedure is commonly called a ‘pseudobulk’ analysis. We used limma to identify differentially expressed genes specific to a given spatial domain (log2FC ≥ 1, multiple-testing adjusted *p* ≤ 0.05) (**Supplementary Table 8**) (56).

Among the four tumor-associated domains, the baseline tumor domain had a small number of differentially expressed genes (*CXCL8*, *MMP12*, *TFF1*, *HBB*). This result indicated limited expression differences in this tumor domain (**Supplementary Table 8**). The intermediate state tumor domain was predominantly present in two CRC-BMets (**Supplementary Fig.10a**). This domain overexpressed histone and HLA family genes, potentially reflecting epigenetic reprogramming and inflammatory processes (**Supplementary Fig.10c, Supplementary Table 8**). Therefore, we focused our analysis on the genomic instability and immune evasive tumor domains; both were consistently represented among all of the metastatic tumors and had significant gene expression differences (**Supplementary Fig. 10a, Supplementary Table 8**).

The genomic instability tumor spatial domain overexpressed genes and pathways associated with increased DNA replication indicating genomic instability, cell proliferation and CRC drivers (e.g., *TSPAN8*) associated with aggressive CRCs (**Fig. 5d, Supplementary Fig. 11a**). In contrast, the immune-evasive tumor domain overexpressed adhesion and differentiation genes and pathways (e.g., *CEACAM6*, *LCN2* and multiple *S100A* family genes) (**Fig. 5e, Supplementary Fig. 10b**). As shown previously (**Fig. 5c**), this domain demonstrated an expression signature indicating immunosuppressive pathways in the brain TME.

Notably, the genomic instability tumor domain had a significantly higher proportion of endothelial cells (Wilcoxon test, *p*=3.9e-12) (**Supplementary Fig. 11c**). The immune evasive tumor domain had significantly higher macrophages (Wilcoxon test, *p*=2.3e-15). Hence, endothelial proximity to tumor cells is associated with a proliferative state of the metastatic CRCs.

### Stromal and macrophage domains mediate angiogenesis and immunosuppression

The stromal-macrophage spatial domain consisted of fibroblasts, macrophage and endothelial cells (**Fig. 5a**). This domain had upregulation of angiogenesis, pro-metastasis and avoiding immune destruction pathways (**Fig. 5c**). Differential expression analysis revealed high expression of ECM related genes and pathways including multiple collagens (*COL1A1*, *COL1A2*, *COL3A1*) and remodeling enzymes (*MMP7*, *SERPINE1*) (**Fig. 5f, Supplementary Fig. 11d, Supplementary Table 8**). This domain overexpressed not only endothelial lineage markers (*PLVAP*) but also genes involved in angiogenesis (*ECSCR*, *VWF*, *FSTL1*). The cytokines *TGFB1* and *TGFB3* were significantly increased in expression, consistent with the features of an immunosuppressive microenvironment. This domain’s macrophages were immunosuppressive TAMs with high expression of *SPP1*, *CD163*, *APOE* and *C1QC*. High expression of *IL2RA* indicated a potential presence of Tregs in this vascularized stroma.

### Adjacent brain parenchyma supports metastatic tumor growth

As noted previously, there were three domains with abundant brain parenchymal cell types. They included: i) protumor brain parenchyma; ii) adjacent brain parenchyma; iii) adjacent microglia brain parenchyma. Differentially expressed genes from all three domains were enriched for pathways related to specific neuronal receptors and synaptic events (**Supplementary Fig. 12a-c**). The protumor brain domain showed upregulated gene expression programs for astrocytic and glial activation, neuronal plasticity and oligodendrocyte/myelin programs (**Supplementary Fig. 12d, Supplementary Table 8**). This domain also expressed angiogenic and inflammatory signatures together with matrix and perivascular cues (*SPP1*, *TGFB2, SPARC*) consistent with a tumor-supportive state. The microglia-enriched brain parenchymal domain showed expression of T-cell activation and checkpoint genes (*GZMK, CTLA4, TIGIT*) indicating local immune activity (**Supplementary Table 8**). In contrast, the adjacent brain parenchymal domain was enriched for homeostatic neuronal and myelination programs with tight-barrier endothelium and normal microglia (*P2RY12*). Overall, this domain lacked pro-tumor signatures.

Each spatial domain had distinct activation of immune pathways (**Supplementary Fig. 12e**). The immunosuppressive stromal–macrophage domain had the highest levels of Treg and exhaustion expression signatures (ANOVA with post-hoc Tukey HSD *p* < 0.05), followed by the protumor brain parenchymal domain. This result indicates their shared role in establishing immunosuppression in the metastatic niche. Cytotoxic effector programs were diminished relative to adjacent brain parenchyma. Within tumor domains, the genomically unstable domain retained high TCR and BCR signaling, suggesting that chromosomal instability may coexist with ongoing immune recognition (57). In contrast, immune-evasive tumor domains upregulated anti-inflammatory T-cell programs. Together, these findings indicate that CRC-BMets establish a vascularized, ECM-rich stroma with abundant macrophages – these cells can contribute to tumor proliferation and immunosuppression within the metastatic brain niche.

### Radiation resistant CRC brain metastases show tumor plasticity and an immunosuppressive TME

Most brain metastasis from epithelial cancers such as non-small cell lung cancer and breast are highly responsive to radiation treatment. However, CRC-BMets are generally refractory to radiation (5). We examined the mechanisms of CRC-BMet resistance to radiotherapy. For this analysis we had five patients (P9, P28, P33, P34, P38) with resected CRC-BMet tissue prior to and then after targeted radiation therapy to the metastatic site. Three patients (P9, P28, P33) represented local recurrence at the same site, while two (P33, P38) recurred at different locations (**Supplementary Table S2**). Illustrating an example, patient P9’s magnetic resonance imaging (**MRI**) showed a local recurrence measuring 2.7 cm along its longest axis, comparable in size to the pre-treatment lesion of 2.0 cm (**Fig. 6a**). These longitudinally matched tissue samples had spatial gene expression data (Visium). Using the results from spatial regions with 100% tumor, we performed differential gene expression with MAST (log2FC ≥ 0.5 and multiple-testing adjusted *p* ≤ 0.05) (32). Patient assignment was used as a random effect to account for inter-patient variability in the mixed effects model, while the number of detected genes per spot was included as a covariate to control for sequencing depth. This procedure provided differential gene expression that was present across all five longitudinal tumor pairs. For all of these BR-MEts the tumor regions had upregulated mesenchymal remodeling genes (*COL1A1*, *COL1A2*, *SPARC*, *KLK6*, *KLK10*, *MMP7*) (**Fig. 6b**, **Supplementary Fig. 13a**) and an increase in EMT signature (**Supplementary Fig. 13c**) (t-test p ≤ 2.83e-09). We used IHC to confirm COL1A1 expression in tumor cells from the same tissues (**Supplementary Fig. 13b**).

**Figure 6:**
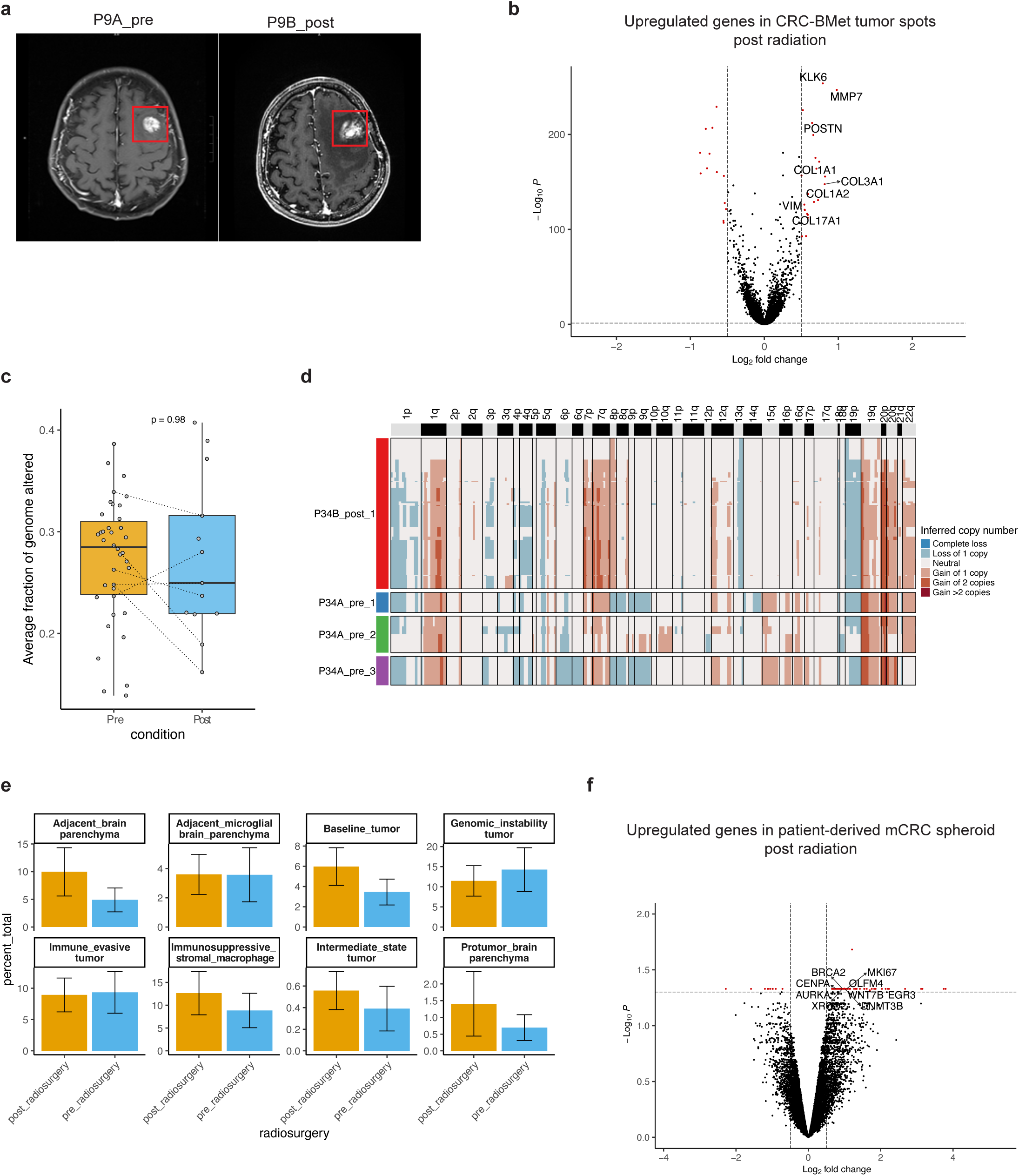
(a) MRI images from P9 depicting longitudinal lesions before and after radiation treatment. (b) Volcano plot highlighting upregulated genes in pure tumor spots from CRC-Bmets longitudinal samples post-radiation compared with pre-radiation. (c) Average inferred fraction of genome altered in pure tumor spots from CRC-BMets post or pre-radiosurgery. Dotted lines connect paired samples from the same patient. (d) Inferred CNV profiles of tumor spots from longitudinal lesions from P34, with suffix indicating subclone. (e) Comparison of proportions of spatial domains between pre and post radiation samples. All comparisons have adjusted *p*□>□0.05 using Wilcoxon test. (f) Volcano plot highlighting upregulated genes in patient-derived CRC-BMet spheroid after radiation treatment.

Next, we compared the CRC-BMets’ chromosomal imbalance features and CIN levels before and after treatment. For all patients the post-radiation treated tumor cells retained the same features as the pre-treatment including chromosomal arm imbalance profiles, clonal composition and levels of genomic instability as indicated by inferred FGA (t-test *p*=0.98) (**Fig. 6c-d, Supplementary Fig. 13 d-g**).

We examined the effect of radiation on cellular organization of the metastatic niche. The proportions of spatial domains were the same when comparing pre- and post-treatment brain metastases (Wilcoxon test, *p*=0.29-0.88) (**Fig. 6e**). Similarly, the proportion of individual cell types (**Supplementary Fig. 13h**) (Wilcoxon signed rank test, multiple-testing adjusted *p* = 0.7-1) also remained the same. Hence, radiation therapy did not change the immunosuppressive stromal-macrophage cellular neighborhoods that surround tumor cells.

To study the acute response of CRC-BMets to radiation, we conducted experiments using a metastatic tumor cell spheroid derived from a patient with CRC brain metastasis (**Methods**). We irradiated the tumor cells with 6 Gy of radiation, harvested the cells after 72 hours and conducted RNA-seq on both the irradiated and non-irradiated cells. To compare the two conditions, we performed differential expression analysis using the limma–voom pipeline with quality weights. This method models the mean–variance relationship of gene expression and assigns weights to account for differences in sequencing depth, allowing the detection of consistent expression changes across replicates (multiple-testing adjusted *p* < 0.05, log2FC > 0.5).

As a result of radiation exposure, the tumor cells had higher expression of DNA repair genes, indicating activation of the DNA damage response after radiation exposure (**Fig. 6f, Supplementary Table 9**). Moreover, with radiation exposure cells simultaneously re-entered the cell cycle as noted by the increased expression of genes that denoted proliferation (*MKI67*, *KIF2C*, *AURKB*). We also observed increased gene expression programs indicating stem cellular processes and cellular plasticity (*OLFM4*, *WNT7B*, *DNMT3B*, *EGR3*, *EGR4*, and *MYBL2*). Overall, these results suggest that CRC-BMets rapidly adapt to radiation using a coordinated increase in DNA repair, proliferation and stem cell-associated regeneration, potentially priming them for the mesenchymal transition observed in patient tumors post-radiation.

### Tumor and macrophage intercellular signaling in the brain TME

We determined which signaling factors mediate communication and response among the adjacent stromal, macrophage and tumor cells in the CNS metastatic niche. This analysis used the CellChat algorithm and the spatial gene expression data (Visium) to identify significant ligand–receptor interactions among spatially proximal cells (58). This program quantifies the average ligand-receptor expression between two cell types as a simplified mass action model followed by permutation for significance testing. It has been widely used for studying signaling pairs in spatial data (59, 60).

Based on the data from adjacent regional spots, we identified 62 significant ligand-receptor pairs (*p* < 0.01) that were consistently detected in at least 80% of CRC-BMets samples (**Supplementary Table 10**). This included CXCL12–CXCR4 signaling between fibroblasts and macrophages and PDGFB–PDGFRB signaling between endothelial cells and fibroblasts. These receptor-ligand pairs are well-established canonical pathways that mediate communication between these cell types and provided an internal validation of this approach (61, 62).

To identify intercellular communication that may influence transcriptional programs in the metastatic TME, we prioritized ligands produced by one cell type that can engage multiple neighboring cell types. We identified five such ligands (**Fig. 7a**). Macrophages had high gene expression of *SPP1*. The complementary integrin or CD44 receptors, both of which bind to SPP1, also had significantly high expression among the adjacent macrophages themselves, as well as with tumor, fibroblast, endothelial, T and NK cells (**Fig. 7b-c**). SPP1 interactions with its receptors drive ECM remodeling, angiogenesis and immunosuppression among these different cell types (63, 64).

**Figure 7:**
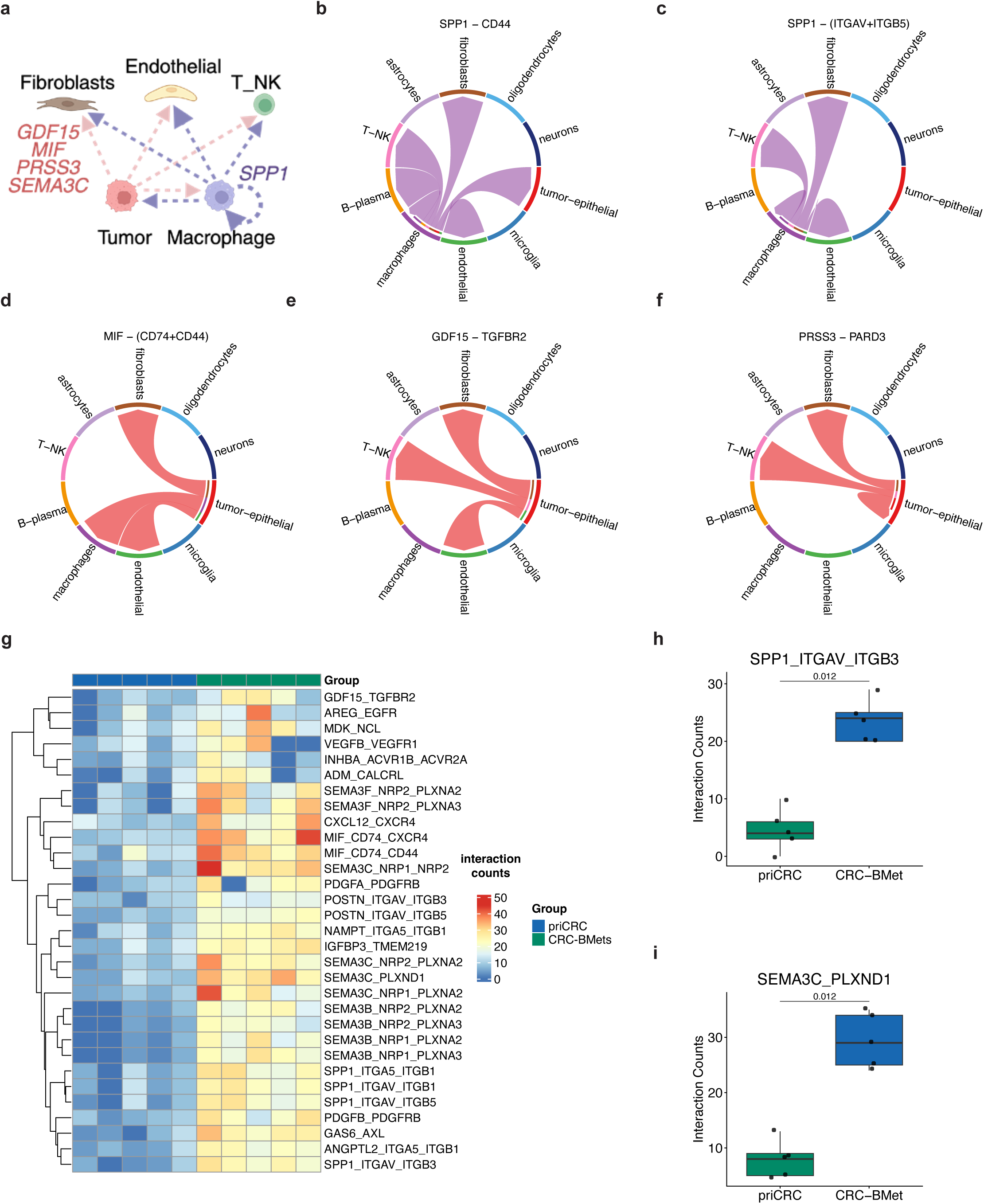
(a) Schematic representation of multi-cellular signaling hubs with tumor or macrophage derived ligands and interacting cell types. (b-f) Chord diagrams indicating cell-cell communication from respective ligands and receptors in P2. Outer bars represent different cell types. Thin inner bars represent target cell types that receive ligand signal from cell type represented by corresponding outer bar. (g-i) Differentially expressed ligand-receptor interaction counts between paired primary CRC and CRC-BMets.

The tumor cells expressed four ligands which can interact with multiple surrounding cell types. For example, the *MIF* ligand that binds to the CD74–CXCR4 and CD44 receptors on fibroblasts, macrophage, and endothelial cells has potential roles in macrophage recruitment, fibrotic remodeling and inflammatory signaling (65, 66) (**Fig. 7d**). The *GDF15* gene’s protein ligand binds to TGFBR2 (**Fig. 7e**) and can activate fibroblasts, promote vascular proliferation and suppress immunity (67–69). *PRSS3* is a ligand gene which acts through stromal and tumor receptors, is linked to metastasis (70) and fibroblast activation (71) (**Fig. 7f**). Finally, *SEMA3C* is a ligand which acts through neuropilin and plexin receptors on endothelial cells and fibroblasts can enhance angiogenesis and fibrosis (72, 73).

To determine whether these ligand-receptor interactions were specific to brain metastasis, we identified the significant ligand–receptor interactions among the five paired primary CRC and CRC-BMets. This analysis was conducted with limma using the interaction counts predicted by CellChat (**Methods**). We identified 28 ligand–receptor pairs with significantly higher interaction counts in the CRC-BMets compared to the primary CRCs (**Fig. 7g**). Notably, *SPP1*, *MIF*, *GDF15* and the *SEMA* family ligand interactions were significantly higher in brain metastases compared to the primary CRCs (**Fig. 7g-i**). Together, these results revealed specific tumor and macrophage ligands that act as multicellular signaling hubs in the brain metastatic TME. Among the neighboring cells, these ligands increase angiogenesis, fibroblast activation and immunosuppression.

### SPP1 regulates macrophage polarization and sustains TME cell interactions

We directly tested the role of *SPP1* in mediating interactions among some of the cell components of the metastatic niche. These experiments involved an in vitro three-dimensional co-culture model to study TME cell interactions (74, 75). The co-cultures had three different cell types including macrophages, fibroblasts and CRC-BMet epithelial cells growing as spheroids. Fibroblasts were represented by the CCD18Co cell line, and macrophages by U937 monocyte-derived macrophages, both of which have been successfully used in TME co-culture models (74, 76, 77). We used CRISPR-Cas9 to introduce a *SPP1* knockout (**KO**) in U937 monocyte cells. The KO relied on two different gRNA constructs (SPP1_KO_1, SPP1_KO_2) for the *SPP1* KO as well as a gRNA control as a *SPP1* wild type (**WT**) control (78). After the CRISPR transduction, we isolated single cells and confirmed the deletion genotype in the *SPP1* KO cells, ensuring that the KO was exclusively represented in the cells (**Supplementary Fig. 14a-b**). Then, the KO and WT monocyte cultures were exposed to phorbol 12-myristate 13-acetate (**PMA**) causing them to differentiate into macrophages (**Methods**). Afterwards, the *SPP1* WT or KO macrophages were grown with the CRC-BMet spheroids and fibroblasts. After 72 hours of incubation, we harvested the cells and then performed scRNA-seq on both the WT and KO co-cultures (**Supplementary Table 11**). We aggregated the data, performed dimensionality reduction and clustering. Relying on canonical marker genes, we identified the three individual cell types among the harvested co-cultures (**Fig. 8a-c**). We characterized the gene expression among the WT macrophages, tumor cells and fibroblasts (Wilcoxon, log2FC= 0.5, adjusted p <= 0.05). Notably, the WT macrophages had increased *SPP1* expression (**Fig.8d**). Tumor cells differentially expressed epithelial genes including *TFF3* and *TSPAN8* – these same genes were also identified among the CRC-BMets tumor epithelium per our spatial analysis. The fibroblasts highly expressed ECM genes including *DCN* and *LOX*. Also, we compared the scRNA-seq data from the co-culture cells with the WT macrophages to the spatial gene expression data from the CRC-BMets. This analysis used the spatial gene expression data (Visium) from regions with 100% fractional representation of each cell type: tumor cells, macrophages or fibroblasts. All three cell types had high concordance in gene expression (average Spearman rho = 0.73, *p* < 2.2e-16) (**Supplementary Fig. 14c**). Overall, the co-culture model recapitulated the cellular gene expression programs identified among the patient metastatic lesions.

**Figure 8:**
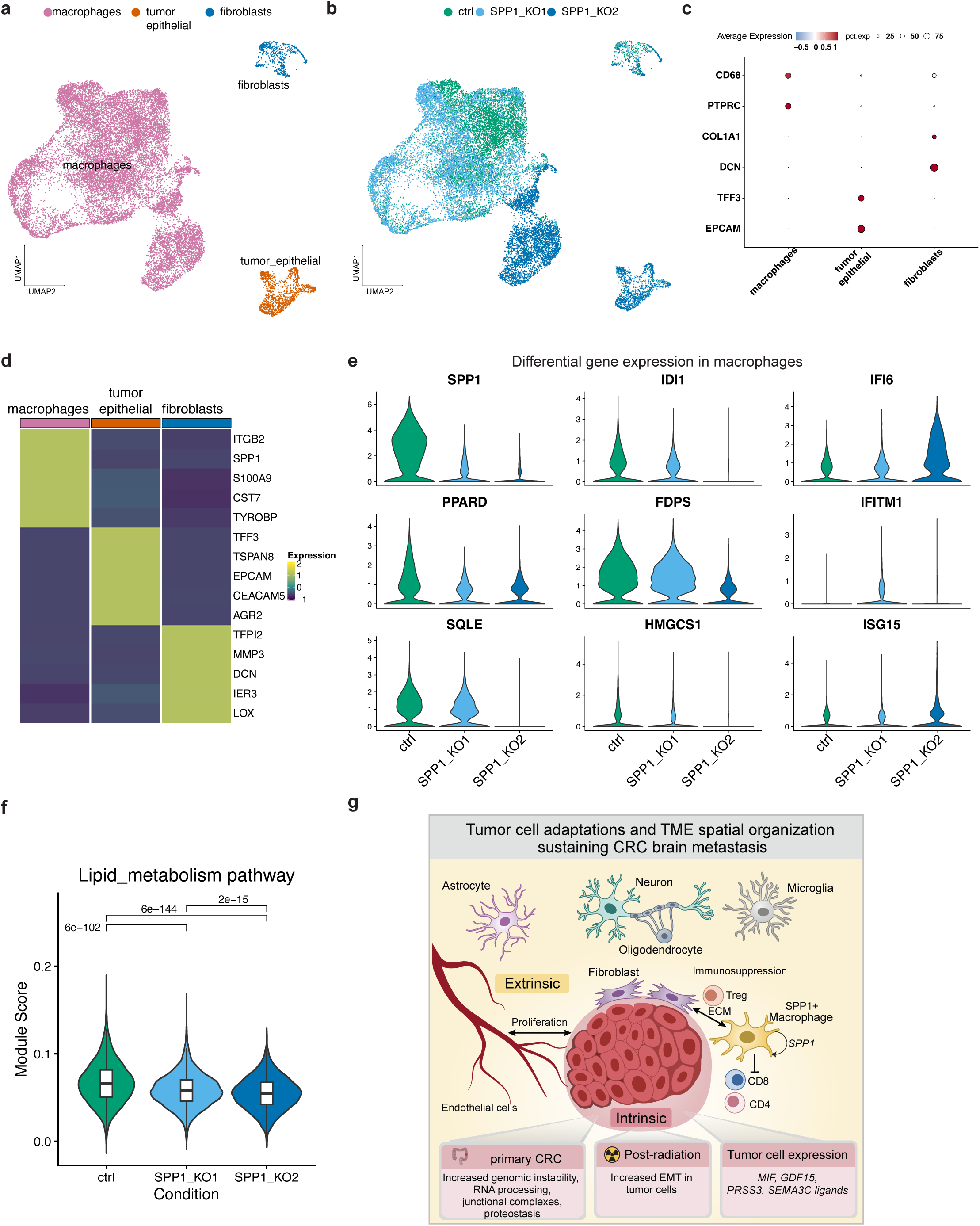
(a-b) UMAP representation of co-culture scRNA-seq colored by (a) cell types and (b) experimental condition. (c) Expression of lineage marker genes in respective cell lineages. Size of the dot is proportional to the percentage of cells expressing the gene. (d) Average scaled expression of top five differentially expressed genes in respective cell lineages from WT co-culture. (e) Violin plots depicting expression of differentially expressed genes in macrophages across co-culture conditions. (f) Lipid metabolism gene signature score in macrophages across co-culture conditions with Kruskal-Wallis *p* following Dunn’s correction. (g) Schematic representation of study findings.

Next, we examined the effects of *SPP1* KO versus the WT macrophages in the co-cultures. We conducted differential expression analysis (Wilcoxon test, log2FC= 0.5, multiple-testing adjusted p ≤ 0.05) between each cell type. In macrophages, *SPP1* was highly downregulated in KO cells as one would expect (**Fig. 8e, Supplementary Table 12**). Also, we observed a significant reduction in expression of genes related to lipid and cholesterol biosynthesis (e.g., *IDI1*, *FDPS*, *SQLE*). The *SPP1* KO cells also had reduced *TREM1* expression, which is associated with a protumor TAM phenotype (79). With both gRNA constructs, we confirmed a significant reduction in lipid metabolism pathway expression in the SPP1 KO compared to the WT macrophages (**Fig. 8f**). Conversely, *SPP1* KO cells had upregulated expression of interferon signaling genes including *IFITM1*, *IFI6* and *ISG15*. Pro-tumor, immunosuppressive TAMs have elevated macrophage lipid metabolism (80). The *SPP1* KO macrophage reduced lipid metabolism pathways and initiated an early shift toward an inflammatory phenotype. These results indicate that *SPP1* acts as an autocrine ligand to sustain macrophage polarization toward a lipid-reprogrammed, immunosuppressive state.

In tumor epithelial cells, the *SPP1* KO reduced the expression of mesenchymal and metastasis associated genes (*VIM*, *S100A family*, *TSPAN1*) while increasing ribosomal and metabolic genes (**Supplementary Fig. 9d, Supplementary Table 13**). There was also upregulated expression of *SRRM2* and *CDKN1C*, linked to RNA processing and cell-cycle regulation respectively. In fibroblasts, the *SPP1* KO led to upregulated gene expression for mesenchymal differentiation (*PDGFRA*, *SOX5*) and phosphatidylinositol signaling (*CDC42BPA*, *PDE1C*, *PLA2G4A*) (**Supplementary Fig. 9e, Supplementary Table 14**).

## DISCUSSION

This study represents the first and most extensive spatial characterization of CRC-Bmets. We identified both extrinsic factors reflecting the surrounding cells in the brain TME and intrinsic factors related to genomic properties of the tumor cells (**Fig. 8g**). Across all metastatic CRCs, there were specific patterns of spatial cellular organization and intercellular signaling. Together, both of these factors contribute to CRC-BMet growth.

One of the most prominent extrinsic cell factors was the tumor epithelium-fibroblast–macrophage cellular hub. A prominent featured embedded in the metastatic TME, these cells jointly support tumor growth via angiogenesis, ECM remodeling, and immunosuppression. We previously identified a similar immunosuppressive macrophage–fibroblast interaction in CRC metastasis to the liver (25). Our results point to the fibroblast-macrophage domain being a conserved tumor-extrinsic program that supports metastatic growth across different organs. SPP1, the macrophage-derived ligand, emerged as a functional regulator of this interaction. We conducted experiments that confirmed that SPP1 can act in an autocrine manner to sustain lipid metabolic reprogramming and immune suppression in macrophages, while influencing fibroblast and tumor compartments through paracrine signaling. Together, these results establish *SPP1* as a critical regulator of macrophage function and a key organizer of the tumor-promoting microenvironment in CRC brain metastasis.

These interacting fibroblasts and macrophages are present across different metastatic sites that include the brain, liver and others. This observation suggests that these TME cells arise from conserved cellular precursors. SPP1+ macrophages in the brain can derive from infiltrating peripheral monocytes, CNS–associated macrophages or resident microglia (21, 81). In the case of brain metastases, recent studies support their origin from peripheral monocytes (26, 27). Potential sources of fibroblasts in CRC-BMets include CNS-resident perivascular or meningeal fibroblasts, pericytes, or bone-marrow–derived mesenchymal fibrocytes (82–84). Together, these cell types form an immunosuppressive stroma that promotes tumor survival and adaptation within the brain niche.

The CRC epithelial tumor cells showed significant expression of ligand genes such as *MIF, GDF15*, *PRSS3*, *SEMA3C.* These ligands may play an important role in establishing metastatic CRC cells in the TME. Future studies will investigate the capacity of these ligands to disrupt multicellular signaling in the TME. Tregs represent another component of the immunosuppressive niche, and their selective modulation or depletion (85) could further enhance antitumor immunity. Finally, tumor-induced remodeling of the brain cellular parenchyma adds an additional layer of immune evasion and stromal adaptation, consistent with co-opting neural processes described in small cell lung cancer brain metastasis (86) and extensively studied in glioblastoma (87, 88).

Among all of the CRC-BMets, the tumor cells had elevated levels of CIN compared to the primary site in the colon. Similar results have been seen in other cancer types with brain metastases and these observations suggest that CIN might be a pan-cancer feature of brain metastases (15, 89). In our cohort, CIN levels in CRC-BMets were not only higher than in matched primary CRCs but also higher than in a bladder metastasis from the same patient. A recent study also demonstrated the highest burden of chromosomal imbalances in CRC-BMets compared to primary CRC or metastases to liver or lung (9). There are different mechanisms by which CIN can promote brain metastasis: these include gene dosage effects or modulation of the microenvironment (90). Moreover, CIN CRCs may represent a potential druggable target in CRC-BMets (91).

To our knowledge, this is the first molecular and spatial analysis showing the consequences of radiation therapy in CRC-BMets. After radiation exposure, resistant clones persisted, leading to recurrence of brain metastases. Clinical studies also point to this radiation resistance present in metastatic brain CRC – namely, that surgical resection of CRC-BMets provides better control compared to radiation (5). We identified EMT as an adaptation in tumor cells, leading to radiation resistance (92). Radiation did not lead to any significant spatial cell remodeling, indicating maintenance of the immunosuppressive spatial domains. These findings are in line with recent data showing that brain metastases from breast cancer resist spatial remodeling after radiation exposure, unlike primary brain tumors such as glioblastoma (50).

Limitations of this study include the clinical variation in the presentation of CRC-BMets among these patients, some of which had different treatment regimens. To mitigate this variation, we focused on the most common features across all of the metastases. The sequencing spatial array (Visium), which provides complete transcriptome features, has lower resolution for discriminating specific cell types compared to the spatial in situ assay (Xenium). However, we gained important insights from this spatial transcriptomics results that included spatial CNV assessment, which has been shown to be robust in a prior study (42). Current ligand-receptor analysis algorithms do not account for deconvolved cell type contributions, potentially reducing specificity. Spatial transcriptomics however permits the evaluation of physiologically relevant spatially restricted communication within a spatial constraint, compared to inferences from dissociated scRNA-seq.

Despite these limitations, our findings establish how the spatial cellular topography sustains CRC-BMets. These experiments provide us with an assessment of how these cellular programs and interactions operate within a patient’s metastatic niche, allowing the prioritization of mechanistic studies of cell states, interactions, domains and therapeutic opportunities.

## METHODS

### Study Approval

This study was conducted in compliance with the Helsinki Declaration and approved by the Institutional Review Board (IRB-66050). Written informed consent was obtained from all patients.

### Visium spatial transcriptomics with CytAssist

Visium libraries were prepared using the Visium CytAssist Spatial Gene Expression for FFPE Kit (version 2, 10x Genomics, CA, USA) as per manufacturer’s protocol. Briefly, 5 μm sections were cut from FFPE blocks and placed onto a histology glass slide. Deparaffinization and hematoxylin and eosin staining was performed by the Stanford Human Pathology Core and whole-slide brightfield images were acquired on a Aperio AT2 whole slide scanner (Leica Biosystems Inc., IL, USA). Coverslips were removed by incubating in xylene followed by rapid freezing as per manufacturer’s protocol. Slides were destained using 0.1N HCl and de-crosslinked, hybridized with human whole transcriptome probes. Following probe ligation, probes were released from the tissue using CytAssist instrument enabled RNA digestion. Probes were captured onto a spatially barcoded Visium CytAssist Spatial Gene Expression v2 slide. Barcoded ligation products were amplified, and sample index PCR was performed to generate sequencing libraries. Cycles for sample index PCR were determined using qPCR as outlined in the manufacturer’s protocol. Image alignment between the whole-slide H&E image and CytAssist image was performed using 10X Genomics Loupe browser (version 6.5.0)

### Visium spatial transcriptomics with direct placement

Sequencing libraries for six samples (P40, P41A, P42, P43, P44, P3) were prepared using the direct placement workflow for the Visium Spatial FFPE Gene Expression Kit (version 1) (10X Genomics). All steps were performed according to the manufacturer’s protocol. Briefly, 5 µm thick tissue sections from the FFPE block were placed onto a Visium Spatial Gene Expression slide using a microtome. Following deparaffinization, hematoxylin and eosin staining was performed, coverslip was mounted using 85% glycerol, and slide was imaged using a Leica DMI 6000 microscope. Slide was subjected to decover slipping and probe hybridization using the Visium Human Transcriptome Probe Set. Sequencing library was constructed following probe ligation, extension and release as per protocol. Cycles for sample index PCR were determined using qPCR as outlined in the manufacturer’s protocol.

### Spatial transcriptomics sequencing data processing

Spatial transcriptomics libraries were sequenced on Illumina NovaSeq sequencers (Illumina, San Diego, CA). Space Ranger version 2.0.1 was used to generate fastq, align reads to GRCh38 together with image alignment, tissue detection, barcode and UMI counting, and generation of a feature-barcode matrix. We constructed Seurat objects from each individual dataset using Seurat (version 4.0.3) with the spot level expression data and corresponding image. We removed spots that expressed fewer than 200 genes. We normalized data using ‘SCTransform’ and used first 20 principal components with a resolution of 0.8 for clustering. For CytAssist datasets, we removed spots with greater than or equal to 10% mitochondrial genes. For direct placement samples, we filtered clusters lacking expression of lineage marker genes. A board-certified pathologist verified that these spots localized to necrotic regions by examining the corresponding H&E image.

### Spatial in situ gene expression imaging

The Xenium assay was used for in situ gene expression with cell segmentation staining (version 1) (10X Genomics). Subset of samples was selected for Xenium based on tissue size. The entire tissue region of these samples was covered by the capture area of each Xenium slide. All steps were performed according to the manufacturer’s protocol. Briefly, 5 µm thick tissue sections from the FFPE block were placed onto a Xenium slide using a microtome. Following deparaffinization, tissues were rehydrated and de-crosslinked to release RNA in situ. Pre-designed Human Immuno-oncology panel together with add-on custom probes (**Supplementary Table 3**) were then added to the tissue for overnight hybridization. Probe ends were ligated to generate a circular DNA probe, which was enzymatically amplified. Slides were incubated overnight with cell segmentation reagents to stain cell membranes and the interior. Following autofluorescence quenching, nuclei were stained with DAPI. Slides were then loaded into the Xenium Analyzer for imaging and decoding. After run completion, hematoxylin and eosin staining was performed as per manufacturer’s protocol. Slides were scanned using an Aperio AT2 whole slide scanner.

Xenium on board analysis was performed using Xenium Analyzer version 2.0. This included cell segmentation, transcript imaging and decoding to generate a cell-feature matrix. We constructed Seurat objects from each individual dataset using Seurat (version 5) (93) with the single-cell gene expression data, the centroid and boundary of each cell and the location of each individual detected transcript. From each individual dataset, we removed cells that expressed fewer than four genes and ten transcripts. We normalized transcript counts to the logarithmic scale and used first 20 principal components with a resolution of 1 for clustering.

These datasets were merged and integrated in Seurat using Harmony [14] (version 0.1.0) with a cluster resolution of 0.8 and the first 20 principal components. From the harmonized object, we identified tumor epithelial cells and non-tumor cells based on marker gene expression. Non-tumor cells were re-clustered with Harmony using a clustering resolution of 0.6. Major TME clusters consisting of lymphocytes, myeloid and stromal or brain parenchyma were identified. These lineages were re-clustered with Harmony to obtain final cell type assignments. At each clustering stage, we filtered clusters lacking expression of lineage marker genes or co-expressing markers of multiple lineages. A pathologist verified that these spots localized to necrotic regions or tissue artefacts respectively by examining the corresponding H&E image.

### Single cell gene expression reference

To perform spot deconvolution, we first assembled an scRNA-seq reference atlas using data generated for this manuscript and publicly available data. Individual datasets were processed using Seurat (version 4) (93).

1. We generated scRNA-seq data from a fresh surgical resection of a CRC brain metastasis (P41A). We filtered genes expressed in less than three cells. We included cells expressing between 200 to 7000 genes, a maximum mitochondrial percentage of 30 and those identified as singlets with DoubletFinder (version 2.0.3).
2. We utilized a CRC brain metastasis patient sample dataset from Gonzalez et. al. (12) obtained from Gene Expression Omnibus GSE186344. Genes and cells were filtered as described above.
3. We included colorectal cancer scRNA-seq from 63 patients obtained from Joanito et. al. (94) using tumor cells identified from metadata labels ‘iCMS2’ and ‘iCMS3’ by the authors. We randomly down sampled this dataset to contain a maximum of 3000 cells per cell type as defined by the original publication.
4. Brain parenchymal cells data from Welch et. al. (95) was obtained from the Spatio-Temporal cell atlas of the human brain (96). The assembled reference was randomly down sampled to include a maximum of 3000 cells from each cell type.

These datasets were merged and integrated in Seurat using the Harmony algorithm [14]. Reference dataset and binned mitochondrial read percentage (with breaks of 10) were provided as the grouping variables in the ‘RunHarmony’ function, and this reduction was used in both ‘RunUMAP’ and ‘FindNeighbors’ functions for clustering. First 20 principal components and a resolution of 1 was used for clustering. Cell lineages were identified based on marker gene expression and confirmed with author annotations for the public datasets. We filtered clusters with mixed cell lineages or ambiguous marker expression, and rare populations from the primary CRC dataset (enteric glial, mast and plasmacytoid dendritic cells) (13% of total cells). The final assembled atlas served as the scRNA-seq reference atlas for CRC-Bmets. For primary CRC atlas, we excluded brain lineages including neurons, astrocytes, oligodendrocytes and microglia.

### Spatial gene expression and cell type deconvolution

We randomly down sampled the scRNA-seq reference datasets by selecting an approximately equal number of cells per lineage, with the total cell count rounded to 10,000. For each CRC-BMet or primary CRC spatial transcriptomics (ST) dataset, we used the corresponding CRC-BMet or primary CRC scRNA-seq reference dataset as input to Cytospace (version 1.0.6) (31) for spot deconvolution. Default options were used with ‘lap_CSPR’ as the solver parameter. Following deconvolution, estimated cellular fractions for each spot in the spatial transcriptomics dataset were obtained.

### Single-cell RNA sequencing

Tissue from P41A CRC-BMet was collected in plain RPMI on ice immediately after resection and dissected with iris scissors. Tissue dissociation was conducted using a combination of enzymatic and mechanical dissociation using a gentleMACS Octo Dissociator (Miltenyi Biotec) as described previously (97). Cells were cryofrozen using 10% DMSO in 90% FBS (ThermoFisher Scientific, Waltham, MA) in a CoolCell freezing container (Larkspur, CA) at -80 °C for 24-72 hours followed by storage in liquid nitrogen. For scRNA-seq, cryofrozen cells were rapidly thawed in a bead bath at 37 °C, washed twice in RPMI + 10% FBS, and filtered successively through 70 μm and 40 μm filters (Flowmi, Bel-Art SP Scienceware, Wayne, NJ). . From the in vitro co-cultures, cell pellet after thawing and washing was treated with TypLE for two minutes at room temperature to improve the generation of a single cell suspension. Live cell counts were obtained using 1:1 trypan blue dilution. Cells were concentrated between 500-1500 live cells/μl.

The scRNA-seq libraries were generated from cell suspensions using Chromium Next GEM Single Cell 5’ version 2 (10X Genomics, Pleasanton, CA, USA) as per manufacturer’s protocol. Ten thousand cells were targeted with 14 PCR cycles for cDNA and library amplification from P41A CRC-BMet. Eight thousand cells were targeted with 11 PCR cycles for cDNA amplification and 14 cycles for library amplification from in vitro co-cultures. A 1% or 2% E-Gel (ThermoFisher Scientific, Waltham, MA, USA) was used for quality control evaluation of intermediate products and sequencing libraries. Qubit (Thermo Fisher Scientific) was used to quantify the libraries as per the manufacturer’s protocol. Libraries were sequenced on Illumina Novaseq sequencers (Illumina, San Diego, CA). Cell Ranger (10x Genomics) ‘mkfastq’ command was used to generate Fastq files. Cell Ranger ‘count’ (version 3.1 for P41A and version 8 for co-culture) was used with default parameters and alignment to GRCh38 to generate a matrix of unique molecular identifier **(UMI)** counts per gene and associated cell barcode.

### RNA and DNA extraction

RNA and DNA extraction from FFPE scrolls was performed using Qiagen AllPrep DNA/RNA FFPE kit or the MagMAX™ FFPE DNA/RNA Ultra Kit using the Kingfisher instrument. DV200 values for RNA were obtained using Agilent Bioanalyzer.

### Whole genome sequencing

WGS from FFPE samples was performed as described previously (98). Briefly, sequencing libraries were prepared using 50 ng of input DNA with the KAPA HyperPlus kit as per the manufacturer’s protocol. Libraries were sequenced on Illumina Novaseq. The sequence data was aligned with BWA using default parameters, against GRCh38 with bin size 250 kb. Since no normal samples were available as a reference, a flat reference with equal coverage per 250 kb bin was created as per CNVkit documentation. Copy number calls were then generated for each tumor sample in comparison to this reference.

### RNA sequencing

RNA was extracted from frozen cell pellet samples using the RNeasy Mini kit. The extracted RNA samples were quantified using a Qubit Flex Fluorometer and their quality was assessed using an Agilent TapeStation 4200. The libraries were generated using the Illumina TruSeq Stranded mRNA kit, following the manufacturer’s protocol with 300 ng RNA input. The process involved poly-A containing mRNA purification using poly-T oligo attached magnetic beads, followed by fragmentation of the mRNA into small pieces using divalent cations under elevated temperature. First strand cDNA synthesis was performed using reverse transcriptase and random primers, with Actinomycin D included to improve strand specificity. Second strand cDNA synthesis was achieved using DNA Polymerase I and RNase H, replacing dTTP with dUTP in the mix to ensure strand specificity. The 3’ ends of the cDNA fragments were adenylated to prevent self-ligation, and adapters were ligated to prepare the cDNA for hybridization onto a flow cell. The cDNA libraries were enriched with PCR and purified. Samples were sequenced on an Illumina NovaSeq X Plus according to the manufacturer’s recommendations. Reads were aligned to GRCh38 using STAR in one-pass mode followed by HTSeq to generate exonic region gene counts.

### Spatial CNV analysis

We inferred copy number from Visium spots comprising 100% tumor epithelial deconvolution fraction. As a reference control derived from the complete CRC-BMet dataset, we randomly down sampled 500 spots each corresponding to 100% macrophage or fibroblast deconvolution fraction with seed set to 123. Count data was used as input to InferCNV (99). Filtering, normalization and centering by normal gene expression was performed using default parameters. A cut-off of 0.1 was used for the minimum average read counts per gene among reference cells. An additional denoising filter was used with a threshold of 0.2. Copy number variation was predicted using the default six state Hidden Markov Model.

Gene-level inferred CNV were aggregated to the chromosome-arm level. The resulting CNV profiles were clustered using the k-means algorithm, with the optimal number of clusters determined by Gap statistics. To identify subclones with consistent large-scale CNV patterns across autosomes, we merged k-means clusters based on their Jaccard similarity, merging clones with a Jaccard index greater than 0.85. Subclones were visualized using ComplexHeatmap. We calculated the inferred FGA per spot as the proportion of aneuploid segments relative to total number of segments. The overall FGA for each sample was computed as the average inferred FGA across individual spots.

### Cell neighborhood analysis on Xenium data

Cell neighborhood analysis was conducted using imcRtools (version 1.8.0) (48). For each dataset, spatial graph was built with a radius of 65 μm using the ‘expansion’ type. Cell type fraction among the neighbors in this graph was then calculated using the ‘aggregateNeighbors’ function. K-means clustering was performed across all samples. K of 5 was selected based on an elbow plot visualizing a range of k’s from 1 to 20.

### Estimating effects of spatial colocalization in Visium data

To investigate spatial relationships between cell type composition in Visium data, we used MistyR (1.10.0) on each individual dataset. We used the deconvolution fractions as the within spot cell type composition, referred to as an ‘intra’ view in the software. To analyze effects of neighboring spots, we added a ‘juxta’ view using the geometry argument with a 250 μm radius. On both these views, we used the ‘run_misty’ and ‘collect_results’ functions with default parameters.

### Spatially aware clustering on Visium data

We used Banksy (version 1.2.0) (53) to cluster Visium by incorporating feature expression data, an average of the features of its spatial neighbors along with neighborhood feature gradients. For each Visium dataset, we constructed a Seurat object and used the ‘RunBanksy’ function with a lambda of 0.2. Individual objects were merged and PCA was performed on the BANKSY matrix. Both the sample and the Visium assay chemistry (CytAssist or direct placement) were used as variables for the ‘RunHarmony’ function. This reduction was used in both ‘RunUMAP’ and ‘FindNeighbors’ functions for clustering with the first 10 principal components and a resolution of 0.2.

### Intercellular communication analysis

We conducted ligand-receptor analysis on each individual Visium dataset using Cellchat (version 2.1.1) (100). Each spot was assigned a cell type based on its maximum deconvolution fraction. Normalized gene expression matrix was used as input to CellChat with the ‘CellChatDB.human’ secreted signaling receptor-ligand interaction database. Overexpressed ligands, receptors, and interactions were identified using ‘identifyOverExpressedGenes’ and ‘identifyOverExpressedInteractions’ functions. Communication probabilities were computed using the ‘computeCommunProb’ function with the truncated mean method, incorporating spatial proximity within a 250 μm interaction range. Number of interactions were calculated using the ‘aggregateNet’ function. To identify consistently activated cell-cell communication pathways across the patient cohort, we performed a prevalence-based screening of ligand-receptor interactions. We extracted significant ligand-receptor interactions (*p* < 0.01) from each sample and implemented a sample-based screening strategy wherein interactions were ranked according to their prevalence across the cohort. We retained high-confidence interactions that demonstrated statistical significance in at least 80% of samples to ensure robust detection of recurrent signaling pathways while minimizing sample-specific artifacts. To investigate alterations in intercellular communication networks between matched primary CRC and CRC-BMets, we performed a comparative analysis of ligand-receptor interaction frequencies across disease states. For each high-confidence ligand-receptor pair identified through prevalence screening, we quantified the number of significant sender-receiver cell-type combinations (*p* < 0.05) within each patient sample, generating a sample-by-interaction count matrix. To identify specific ligand-receptor pairs exhibiting differential activation, we employed linear modeling using the limma framework (56) on the scaled interaction count matrix, with disease state (primary or metastasis) as the predictor variable.

### Differential expression analysis

Comparisons between paired or longitudinal samples were conducted using MAST (32) (version 1.18). Genes expressed in more than 10% of cells were included, using log-normalized expression values and modeling patient as a random effect to account for inter-patient variability. The number of detected genes per spot was included as a covariate to control for sequencing depth.

From the spatially aware clustering data, one-vs-rest differential expression analysis was conducted using a pseudobulk approach. Gene counts were aggregated by sample and spatial domain using the Seurat AggregateExpression function. The resulting pseudobulk matrices were normalized using the TMM method, and lowly expressed genes were filtered with filterByExpr from edgeR (version 3.34). Differential expression was performed with the limma–voom pipeline (version 3.48.3) (101). The design matrix included sample as a covariate to account for inter-sample variability and domain as the variable of interest. Genes with significant expression differences (FDR ≤ 0.05) were identified using empirical Bayes moderation.

Bulk RNA-seq differential expression analysis was performed using the same limma–voom framework. Gene counts were filtered to retain genes with at least ten counts in two or more samples and normalized using the TMM method. Because one treatment replicate exhibited a substantially higher library size, the analysis used voomWithQualityWeights to estimate sample-specific precision weights and correct for differences in library composition and sequencing depth. In all other analyses, differential expression was performed using Seurat ‘FindMarkers’ or ‘FindAllMarkers’ functions with minimum of 25% percent expressing cells.

### Gene signature analysis

CIN70 gene signature was obtained from the original publication (46). Hallmark EMT gene set and Reactome metabolism of lipids gene set was obtained from msigdbr (version 7.5.1) (40). Gene signatures were scored per spot using the Seurat ‘AddModuleScore function’. Additionally, we used GSVA (version 1.4) (102) with a Gaussian distribution to score gene sets. This included all hallmark gene sets obtained from msigdbr. Tumor and TME hallmark gene sets were obtained from the original publication (55). Treg and anti-inflammatory gene signatures were obtained from Azizi et. al. (103). Reactome BCR and TCR signaling were obtained from msigdbr. Exhaustion signature was obtained from Zheng et. al. (104). Cytotoxic effectors were compiled from prior publications (105, 106) as we previously described (107).

Overrepresentation pathway analysis was conducted using enrichR (39) (version 3).

### Cell culture

PDM-104, a CRC-BMet patient-derived spheroid cell line generated by the Human Cancer Models Initiative (**HCMI**), was obtained from the American Type Culture Collection (**ATCC**). Cells were maintained in ultra-low attachment flasks in Propagenix conditioned media (catalog number 256-100) supplemented with 9.0 ng/mL cholera toxin (Sigma Aldrich C8052). Cas9 expressing U937 cells were a gift from Dr. Michael Bassik, Stanford University and were maintained in RPMI with 10% FBS. CCD-18Co were obtained from the ATCC and cultured in DMEM with 10% FBS. For co-culture setups, WT or SPP1 KO U937 monocytes were differentiated in ultra-low attachment 96-well plates using 40 ng/ml PMA for 24 hours, followed by culture for 48 hours without PMA. CCD18-Co and PDM-104 cells were added to differentiated U937s in a 2:1:1 macrophage-tumor-fibroblast ratio and all cells were co-cultured for a total of 72 hours. HEK293T cells (ATCC) were maintained in DMEM with 10% FBS.

### Radiation

PDM-104 cells in ultra-low attachment 96 well plates were seeded at 40000 cells per well. 24 hours after seeding, cells were irradiated with 6 Gy. Cells were cultured for 72 hours in three independent experiments.

### SPP1 KO

gRNAs targeting SPP1 or safe targeting gRNA (**Supplementary Table 15**) were cloned into pMCB320 (Addgene). Plasmids were packaged into lentivirus in HEK293T using pMDLg/pRRE, pCMV-VSV-G and pRSV-Rev (Addgene). Following lentivirus delivery to U937 cells, successfully transduced cells were sorted based on mCherry expression. For SPP1 KO, single cells were deposited into 96-well plates to establish clonal populations. Clones were expanded and genomic DNA was extracted using Qiagen All Prep DNA/RNA kit. SPP1 target regions were PCR-amplified (**Supplementary Table 15**). Amplicons were prepared for sequencing using the Illumina TruSeq DNA library preparation kit and sequenced on the Illumina iSeq platform. Knockout efficiency was assessed using CRISPResso2 (108).

### Immunohistochemistry

IHC was performed as described previously (109). Briefly, FFPE sections were deparaffinized, rehydrated and blocked. Citrate antigen retrieval was used for COL1A1 (catalog number 72026, Cell Signaling Technology), CD79A (catalog number 13333, Cell Signaling Technology) and CD4 (catalog number 48274). EDTA antigen retrieval was used for CD8 (catalog number 70306, Cell Signaling Technology) and CLDN3 (catalog number ab214487, Abcam). Following respective primary and secondary antibody incubation, staining was visualized with DAB.

### Additional analysis and visualization

Additional analysis or visualization was conducted using R packages ggplot2 (version 3.4.2), ggpubr (version 0.4.0), EnhancedVolcano (1.10), pheatmap (1.10.12), ComplexHeatmap (2.9.3), tidyr (1.1.3), broom (0.7.6), dplyr(1.1.4), plyr (1.8.6), rstatix (0.7.0), viridis (0.6.1), pals (1.7) in R version >4.3. Adobe Illustrator (26.4) was used to edit and assemble figures.

## DATA AND CODE AVAILABILITY

Sequencing data has been deposited to dbGAP identifier phs003794. Xenium and Visium matrices is available on Zenodo. Code is available on Zenodo.

## AUTHORS’ CONTRIBUTIONS

AS was involved in conception and design of the study, development of methodology, acquisition of data, analysis and interpretation of data and writing of the manuscript. JIK, RM, HS were involved in the acquisition of data. XB, MZ, SG, AK, SH were involved in the analysis and interpretation of data. ML and AL performed clinical chart review. AL, ML, CP, CJ, JS, HV and MG were involved in the clinical translational component. HPJ oversaw the conception and design of the study, data analysis, interpretation of data and writing of the manuscript.

## ACKNOWLEDGEMENTS

We are grateful to all patients who participated in the study. This work was supported by the US National Institutes of Health grants U54CA261717 and NIH T32 CA009695 (ML). Fig. 7a was created using Biorender.com.

## ADDITIONAL FILES

**Additional file 1:** Supplementary tables S1 – 15. format: XLSX

**Additional file 2:** Supplementary information - supplementary figure legends and supplementary figures S1 – S14, format: PDF

## REFERENCES

1. Qiu M, Hu J, Yang D, Cosgrove DP, Xu R. Pattern of distant metastases in colorectal cancer: a SEER based study. Oncotarget. 2015;6(36):38658–66.

2. Muller S, Kohler F, Hendricks A, Kastner C, Borner K, Diers J, et al. Brain Metastases from Colorectal Cancer: A Systematic Review of the Literature and Meta-Analysis to Establish a Guideline for Daily Treatment. Cancers (Basel). 2021;13(4).

3. Christensen TD, Spindler KL, Palshof JA, Nielsen DL. Systematic review: brain metastases from colorectal cancer--Incidence and patient characteristics. BMC Cancer. 2016;16:260.

4. Frazier JL, Batra S, Kapor S, Vellimana A, Gandhi R, Carson KA, et al. Stereotactic radiosurgery in the management of brain metastases: an institutional retrospective analysis of survival. Int J Radiat Oncol Biol Phys. 2010;76(5):1486–92.

5. Chang Y, Wong CE, Lee PH, Huang CC, Lee JS. Survival Outcome of Surgical Resection vs. Radiotherapy in Brain Metastasis From Colorectal Cancer: A Meta-Analysis. Front Med (Lausanne). 2022;9:768896.

6. Mjahed RB, Astaras C, Roth A, Koessler T. Where Are We Now and Where Might We Be Headed in Understanding and Managing Brain Metastases in Colorectal Cancer Patients? Curr Treat Options Oncol. 2022;23(7):980–1000.

7. Zhang Q, Abdo R, Iosef C, Kaneko T, Cecchini M, Han VK, Li SS. The spatial transcriptomic landscape of non-small cell lung cancer brain metastasis. Nat Commun. 2022;13(1):5983.

8. Meinhardt A, Hedger MP. Immunological, paracrine and endocrine aspects of testicular immune privilege. Mol Cell Endocrinol. 2011;335(1):60–8.

9. Golas MM, Gunawan B, Gutenberg A, Danner BC, Gerdes JS, Stadelmann C, et al. Cytogenetic signatures favoring metastatic organotropism in colorectal cancer. Nat Commun. 2025;16(1):3261.

10. Gandini A, Puglisi S, Pirrone C, Martelli V, Catalano F, Nardin S, et al. The role of immunotherapy in microsatellites stable metastatic colorectal cancer: state of the art and future perspectives. Front Oncol. 2023;13:1161048.

11. Bhambhvani HP, Granucci M, Rodrigues A, Kakusa BW, Hayden Gephart M. The primary sites leading to brain metastases: Shifting trends at a tertiary care center. J Clin Neurosci. 2020;80:121–4.

12. Gonzalez H, Mei W, Robles I, Hagerling C, Allen BM, Hauge Okholm TL, et al. Cellular architecture of human brain metastases. Cell. 2022;185(4):729–45 e20.

13. Tagore S, Caprio L, Amin AD, Bestak K, Luthria K, D’Souza E, et al. Single-cell and spatial genomic landscape of non-small cell lung cancer brain metastases. Nat Med. 2025;31(4):1351–63.

14. Biermann J, Melms JC, Amin AD, Wang Y, Caprio LA, Karz A, et al. Dissecting the treatment-naive ecosystem of human melanoma brain metastasis. Cell. 2022;185(14):2591–608 e30.

15. Xing X, Zhong J, Biermann J, Duan H, Zhang X, Shi Y, et al. Pan-cancer human brain metastases atlas at single-cell resolution. Cancer Cell. 2025.

16. Oliveira MF, Romero JP, Chung M, Williams SR, Gottscho AD, Gupta A, et al. High-definition spatial transcriptomic profiling of immune cell populations in colorectal cancer. Nat Genet. 2025;57(6):1512–23.

17. Su A, Lee H, Tran M, Dela Cruz RC, Sathe A, Bai X, et al. The single-cell spatial landscape of stage III colorectal cancers. NPJ Precis Oncol. 2025;9(1):101.

18. Nieto P, Elosua-Bayes M, Trincado JL, Marchese D, Massoni-Badosa R, Salvany M, et al. A single-cell tumor immune atlas for precision oncology. Genome Res. 2021;31(10):1913–26.

19. Zheng L, Qin S, Si W, Wang A, Xing B, Gao R, et al. Pan-cancer single-cell landscape of tumor-infiltrating T cells. Science. 2021;374(6574):abe6474.

20. Dorrier CE, Jones HE, Pintaric L, Siegenthaler JA, Daneman R. Emerging roles for CNS fibroblasts in health, injury and disease. Nat Rev Neurosci. 2022;23(1):23–34.

21. Sankowski R, Suss P, Benkendorff A, Bottcher C, Fernandez-Zapata C, Chhatbar C, et al. Multiomic spatial landscape of innate immune cells at human central nervous system borders. Nat Med. 2024;30(1):186–98.

22. Shen J, Bian N, Zhao L, Wei J. The role of T-lymphocytes in central nervous system diseases. Brain Res Bull. 2024;209:110904.

23. Palma A. The Landscape of SPP1 (+) Macrophages Across Tissues and Diseases: A Comprehensive Review. Immunology. 2025;176(2):179–96.

24. Millet A, Ledo JH, Tavazoie SF. An exhausted-like microglial population accumulates in aged and APOE4 genotype Alzheimer’s brains. Immunity. 2024;57(1):153–70 e6.

25. Sathe A, Mason K, Grimes SM, Zhou Z, Lau BT, Bai X, et al. Colorectal Cancer Metastases in the Liver Establish Immunosuppressive Spatial Networking between Tumor-Associated SPP1+ Macrophages and Fibroblasts. Clin Cancer Res. 2023;29(1):244–60.

26. Friebel E, Kapolou K, Unger S, Nunez NG, Utz S, Rushing EJ, et al. Single-Cell Mapping of Human Brain Cancer Reveals Tumor-Specific Instruction of Tissue-Invading Leukocytes. Cell. 2020;181(7):1626–42 e20.

27. Bowman RL, Klemm F, Akkari L, Pyonteck SM, Sevenich L, Quail DF, et al. Macrophage Ontogeny Underlies Differences in Tumor-Specific Education in Brain Malignancies. Cell Rep. 2016;17(9):2445–59.

28. Ewing-Crystal NA, Mroz NM, Larpthaveesarp A, Lizama CO, Pennington R, Chiaranunt P, et al. Dynamic fibroblast-immune interactions shape recovery after brain injury. Nature. 2025;646(8086):934–44.

29. Qiu Y, Shen X, Ravid O, Atrakchi D, Rand D, Wight AE, et al. Definition of the contribution of an Osteopontin-producing CD11c(+) microglial subset to Alzheimer’s disease. Proc Natl Acad Sci U S A. 2023;120(6):e2218915120.

30. Laviron M, Boissonnas A. Ontogeny of Tumor-Associated Macrophages. Front Immunol. 2019;10:1799.

31. Vahid MR, Brown EL, Steen CB, Zhang W, Jeon HS, Kang M, et al. High-resolution alignment of single-cell and spatial transcriptomes with CytoSPACE. Nat Biotechnol. 2023;41(11):1543–8.

32. Finak G, McDavid A, Yajima M, Deng J, Gersuk V, Shalek AK, et al. MAST: a flexible statistical framework for assessing transcriptional changes and characterizing heterogeneity in single-cell RNA sequencing data. Genome Biol. 2015;16:278.

33. Chaturvedi P, Belmont AS. Nuclear speckle biology: At the cross-roads of discovery and functional analysis. Curr Opin Cell Biol. 2024;91:102438.

34. Ivanova OM, Anufrieva KS, Kazakova AN, Malyants IK, Shnaider PV, Lukina MM, Shender VO. Non-canonical functions of spliceosome components in cancer progression. Cell Death Dis. 2023;14(2):77.

35. Alexander KA, Yu R, Skuli N, Coffey NJ, Nguyen S, Faunce CL, et al. Nuclear speckles regulate functional programs in cancer. Nat Cell Biol. 2025;27(2):322–35.

36. Kyuno D, Takasawa A, Kikuchi S, Takemasa I, Osanai M, Kojima T. Role of tight junctions in the epithelial-to-mesenchymal transition of cancer cells. Biochim Biophys Acta Biomembr. 2021;1863(3):183503.

37. Ji W, Zhuang X, Jiang WG, Martin TA. Tight junctional protein family, Claudins in cancer and cancer metastasis. Front Oncol. 2025;15:1596460.

38. Hsu SK, Chiu CC, Dahms HU, Chou CK, Cheng CM, Chang WT, et al. Unfolded Protein Response (UPR) in Survival, Dormancy, Immunosuppression, Metastasis, and Treatments of Cancer Cells. Int J Mol Sci. 2019;20(10).

39. Kuleshov MV, Jones MR, Rouillard AD, Fernandez NF, Duan Q, Wang Z, et al. Enrichr: a comprehensive gene set enrichment analysis web server 2016 update. Nucleic acids research. 2016;44(W1):W90–7.

40. Liberzon A, Birger C, Thorvaldsdottir H, Ghandi M, Mesirov JP, Tamayo P. The Molecular Signatures Database (MSigDB) hallmark gene set collection. Cell Syst. 2015;1(6):417–25.

41. Tickle T, Tirosh I, Georgescu C, Brown M, Haas B. inferCNV of the Trinity CTAT Project: Klarman Cell Observatory, Broad Institute of MIT and Harvard, Cambridge, MA, USA; 2019 [Available from: https://github.com/broadinstitute/inferCNV.

42. Erickson A, He M, Berglund E, Marklund M, Mirzazadeh R, Schultz N, et al. Spatially resolved clonal copy number alterations in benign and malignant tissue. Nature. 2022;608(7922):360–7.

43. Wu CY, Rong J, Sathe A, Hess PR, Lau BT, Grimes SM, et al. Cancer subclone detection based on DNA copy number in single-cell and spatial omic sequencing data. Nat Methods. 2025;22(9):1846–56.

44. Bai X, Lau Billy T, Sathe A, Grimes Susan M, Almeda-Nostine A, Ji Hanlee P. Single-cell aneuploidy and chromosomal arm imbalances define subclones with divergent transcriptomic phenotypes. NAR Genomics and Bioinformatics. 2025;7(4).

45. Yukihiro K, Sekino Y, Kobayashi G, Hatayama T, Shikuma H, Tasaka R, et al. Comprehensive Analysis of Fraction of Genome Altered in Prostate Cancer Treatment. Prostate. 2025.

46. Carter SL, Eklund AC, Kohane IS, Harris LN, Szallasi Z. A signature of chromosomal instability inferred from gene expression profiles predicts clinical outcome in multiple human cancers. Nat Genet. 2006;38(9):1043–8.

47. Bhate SS, Barlow GL, Schurch CM, Nolan GP. Tissue schematics map the specialization of immune tissue motifs and their appropriation by tumors. Cell Syst. 2022;13(2):109–30 e6.

48. Windhager J, Zanotelli VRT, Schulz D, Meyer L, Daniel M, Bodenmiller B, Eling N. An end-to-end workflow for multiplexed image processing and analysis. Nat Protoc. 2023;18(11):3565–613.

49. Salie H, Wischer L, D’Alessio A, Godbole I, Suo Y, Otto-Mora P, et al. Spatial single-cell profiling and neighbourhood analysis reveal the determinants of immune architecture connected to checkpoint inhibitor therapy outcome in hepatocellular carcinoma. Gut. 2025;74(3):451–66.

50. Watson SS, Duc B, Kang Z, de Tonnac A, Eling N, Font L, et al. Microenvironmental reorganization in brain tumors following radiotherapy and recurrence revealed by hyperplexed immunofluorescence imaging. Nat Commun. 2024;15(1):3226.

51. Caramello A, Fancy N, Tournerie C, Eklund M, Chau V, Adair E, et al. Intracellular accumulation of amyloid-ss is a marker of selective neuronal vulnerability in Alzheimer’s disease. Nat Commun. 2025;16(1):5189.

52. Tanevski J, Flores ROR, Gabor A, Schapiro D, Saez-Rodriguez J. Explainable multiview framework for dissecting spatial relationships from highly multiplexed data. Genome Biol. 2022;23(1):97.

53. Singhal V, Chou N, Lee J, Yue Y, Liu J, Chock WK, et al. BANKSY unifies cell typing and tissue domain segmentation for scalable spatial omics data analysis. Nat Genet. 2024;56(3):431–41.

54. Ruitenberg MJ, Nguyen QH. Cellular neighborhood analysis in spatial omics reveals new tissue domains and cell subtypes. Nat Genet. 2024;56(3):362–4.

55. Sibai M, Cervilla S, Grases D, Musulen E, Lazcano R, Mo CK, et al. The spatial landscape of cancer hallmarks reveals patterns of tumor ecological dynamics and drug sensitivity. Cell Rep. 2025;44(2):115229.

56. Ritchie ME, Phipson B, Wu D, Hu Y, Law CW, Shi W, Smyth GK. limma powers differential expression analyses for RNA-sequencing and microarray studies. Nucleic acids research. 2015;43(7):e47.

57. Li J, Hubisz MJ, Earlie EM, Duran MA, Hong C, Varela AA, et al. Non-cell-autonomous cancer progression from chromosomal instability. Nature. 2023;620(7976):1080–8.

58. Jin S, Guerrero-Juarez CF, Zhang L, Chang I, Ramos R, Kuan CH, et al. Inference and analysis of cell-cell communication using CellChat. Nat Commun. 2021;12(1):1088.

59. Jimenez-Santos MJ, Garcia-Martin S, Rubio-Fernandez M, Gomez-Lopez G, Al-Shahrour F. Spatial transcriptomics in breast cancer reveals tumour microenvironment-driven drug responses and clonal therapeutic heterogeneity. NAR Cancer. 2024;6(4):zcae046.

60. Reina-Campos M, Monell A, Ferry A, Luna V, Cheung KP, Galletti G, et al. Tissue-resident memory CD8 T cell diversity is spatiotemporally imprinted. Nature. 2025;639(8054):483–92.

61. Wu X, Qian L, Zhao H, Lei W, Liu Y, Xu X, et al. CXCL12/CXCR4: An amazing challenge and opportunity in the fight against fibrosis. Ageing Res Rev. 2023;83:101809.

62. Andrae J, Gallini R, Betsholtz C. Role of platelet-derived growth factors in physiology and medicine. Genes Dev. 2008;22(10):1276–312.

63. Klement JD, Paschall AV, Redd PS, Ibrahim ML, Lu C, Yang D, et al. An osteopontin/CD44 immune checkpoint controls CD8+ T cell activation and tumor immune evasion. J Clin Invest. 2018;128(12):5549–60.

64. Yue B, Xiong D, Chen J, Yang X, Zhao J, Shao J, et al. SPP1 induces idiopathic pulmonary fibrosis and NSCLC progression via the PI3K/Akt/mTOR pathway. Respir Res. 2024;25(1):362.

65. Zhang Y, Lu S, Fan S, Xu L, Jiang X, Wang K, Cai B. Macrophage migration inhibitory factor activates the inflammatory response in joint capsule fibroblasts following post-traumatic joint contracture. Aging (Albany NY). 2021;13(4):5804–23.

66. Sumaiya K, Langford D, Natarajaseenivasan K, Shanmughapriya S. Macrophage migration inhibitory factor (MIF): A multifaceted cytokine regulated by genetic and physiological strategies. Pharmacol Ther. 2022;233:108024.

67. Wang S, Li M, Zhang W, Hua H, Wang N, Zhao J, et al. Growth differentiation factor 15 promotes blood vessel growth by stimulating cell cycle progression in repair of critical-sized calvarial defect. Sci Rep. 2017;7(1):9027.

68. Reyes J, Yap GS. Emerging Roles of Growth Differentiation Factor 15 in Immunoregulation and Pathogenesis. J Immunol. 2023;210(1):5–11.

69. Guo H, Zhao X, Li H, Liu K, Jiang H, Zeng X, et al. GDF15 Promotes Cardiac Fibrosis and Proliferation of Cardiac Fibroblasts via the MAPK/ERK1/2 Pathway after Irradiation in Rats. Radiat Res. 2021;196(2):183–91.

70. Zhou M, Li K, Luo KQ. Shear Stress Drives the Cleavage Activation of Protease-Activated Receptor 2 by PRSS3/Mesotrypsin to Promote Invasion and Metastasis of Circulating Lung Cancer Cells. Adv Sci (Weinh). 2023;10(25):e2301059.

71. Tabib T, Huang M, Morse N, Papazoglou A, Behera R, Jia M, et al. Myofibroblast transcriptome indicates SFRP2(hi) fibroblast progenitors in systemic sclerosis skin. Nat Commun. 2021;12(1):4384.

72. Iragavarapu-Charyulu V, Wojcikiewicz E, Urdaneta A. Semaphorins in Angiogenesis and Autoimmune Diseases: Therapeutic Targets? Front Immunol. 2020;11:346.

73. Hao J, Yu JS. Semaphorin 3C and Its Receptors in Cancer and Cancer Stem-Like Cells. Biomedicines. 2018;6(2).

74. Wallisch S, Neef SK, Denzinger L, Monch D, Koch J, Marzi J, et al. Protocol for establishing a coculture with fibroblasts and colorectal cancer organoids. STAR Protoc. 2023;4(3):102481.

75. Wang J, Tao X, Zhu J, Dai Z, Du Y, Xie Y, et al. Tumor organoid-immune co-culture models: exploring a new perspective of tumor immunity. Cell Death Discov. 2025;11(1):195.

76. Maolake A, Izumi K, Shigehara K, Natsagdorj A, Iwamoto H, Kadomoto S, et al. Tumor-associated macrophages promote prostate cancer migration through activation of the CCL22-CCR4 axis. Oncotarget. 2017;8(6):9739–51.

77. Yu J, Xu Z, Guo J, Yang K, Zheng J, Sun X. Tumor-associated macrophages (TAMs) depend on MMP1 for their cancer-promoting role. Cell Death Discov. 2021;7(1):343.

78. Morgens DW, Wainberg M, Boyle EA, Ursu O, Araya CL, Tsui CK, et al. Genome-scale measurement of off-target activity using Cas9 toxicity in high-throughput screens. Nat Commun. 2017;8:15178.

79. Colonna M. The biology of TREM receptors. Nat Rev Immunol. 2023;23(9):580–94.

80. Qiao X, Hu Z, Xiong F, Yang Y, Peng C, Wang D, Li X. Lipid metabolism reprogramming in tumor-associated macrophages and implications for therapy. Lipids Health Dis. 2023;22(1):45.

81. Schulz M, Michels B, Niesel K, Stein S, Farin H, Rodel F, Sevenich L. Cellular and Molecular Changes of Brain Metastases-Associated Myeloid Cells during Disease Progression and Therapeutic Response. iScience. 2020;23(6):101178.

82. Wang Z, Liu J, Huang H, Ye M, Li X, Wu R, et al. Metastasis-associated fibroblasts: an emerging target for metastatic cancer. Biomark Res. 2021;9(1):47.

83. Sosa MJ, Shih AY, Bonney SK. The elusive brain perivascular fibroblast: a potential role in vascular stability and homeostasis. Front Cardiovasc Med. 2023;10:1283434.

84. Ippolitov D, Arreza L, Munir MN, Hombach-Klonisch S. Brain Microvascular Pericytes-More than Bystanders in Breast Cancer Brain Metastasis. Cells. 2022;11(8).

85. Iglesias-Escudero M, Arias-Gonzalez N, Martinez-Caceres E. Regulatory cells and the effect of cancer immunotherapy. Mol Cancer. 2023;22(1):26.

86. Qu F, Brough SC, Michno W, Madubata CJ, Hartmann GG, Puno A, et al. Crosstalk between small-cell lung cancer cells and astrocytes mimics brain development to promote brain metastasis. Nat Cell Biol. 2023;25(10):1506–19.

87. Taylor KR, Barron T, Hui A, Spitzer A, Yalcin B, Ivec AE, et al. Glioma synapses recruit mechanisms of adaptive plasticity. Nature. 2023;623(7986):366–74.

88. Krishna S, Choudhury A, Keough MB, Seo K, Ni L, Kakaizada S, et al. Glioblastoma remodelling of human neural circuits decreases survival. Nature. 2023;617(7961):599–607.

89. Tagore S, Caprio L, Amin AD, Bestak K, Luthria K, D’Souza E, et al. Single-cell and spatial genomic landscape of non-small cell lung cancer brain metastases. Nat Med. 2025;31(4):1351–63.

90. Li J, Hubisz MJ, Earlie EM, Duran MA, Hong C, Varela AA, et al. Non-cell-autonomous cancer progression from chromosomal instability. Nature. 2023;620(7976):1080–8.

91. Bhatia S, Khanna KK, Duijf PHG. Targeting chromosomal instability and aneuploidy in cancer. Trends Pharmacol Sci. 2024;45(3):210–24.

92. Qiao L, Chen Y, Liang N, Xie J, Deng G, Chen F, et al. Targeting Epithelial-to-Mesenchymal Transition in Radioresistance: Crosslinked Mechanisms and Strategies. Front Oncol. 2022;12:775238.

93. Butler A, Hoffman P, Smibert P, Papalexi E, Satija R. Integrating single-cell transcriptomic data across different conditions, technologies, and species. Nat Biotechnol. 2018;36(5):411–20.

94. Joanito I, Wirapati P, Zhao N, Nawaz Z, Yeo G, Lee F, et al. Single-cell and bulk transcriptome sequencing identifies two epithelial tumor cell states and refines the consensus molecular classification of colorectal cancer. Nat Genet. 2022;54(7):963–75.

95. Welch JD, Kozareva V, Ferreira A, Vanderburg C, Martin C, Macosko EZ. Single-Cell Multi-omic Integration Compares and Contrasts Features of Brain Cell Identity. Cell. 2019;177(7):1873–87 e17.

96. Song L, Pan S, Zhang Z, Jia L, Chen WH, Zhao XM. STAB: a spatio-temporal cell atlas of the human brain. Nucleic Acids Res. 2021;49(D1):D1029–D37.

97. Sathe A, Grimes SM, Lau BT, Chen J, Suarez C, Huang RJ, et al. Single-Cell Genomic Characterization Reveals the Cellular Reprogramming of the Gastric Tumor Microenvironment. Clin Cancer Res. 2020.

98. Shin G, Greer SU, Hopmans E, Grimes SM, Lee H, Zhao L, et al. Profiling diverse sequence tandem repeats in colorectal cancer reveals co-occurrence of microsatellite and chromosomal instability involving Chromosome 8. Genome Med. 2021;13(1):145.

99. Tickle T, Tirosh I, Georgescu C, Brown M, Haas B. inferCNV of the Trinity CTAT Project: Klarman Cell Observatory, Broad Institute of MIT and Harvard, Cambridge, MA, USA; 2019 [Available from: https://github.com/broadinstitute/inferCNV.

100. Jin S, Plikus MV, Nie Q. CellChat for systematic analysis of cell-cell communication from single-cell transcriptomics. Nat Protoc. 2025;20(1):180–219.

101. Law CW, Chen Y, Shi W, Smyth GK. voom: Precision weights unlock linear model analysis tools for RNA-seq read counts. Genome Biol. 2014;15(2):R29.

102. Hanzelmann S, Castelo R, Guinney J. GSVA: gene set variation analysis for microarray and RNA-seq data. BMC Bioinformatics. 2013;14:7.

103. Azizi E, Carr AJ, Plitas G, Cornish AE, Konopacki C, Prabhakaran S, et al. Single-Cell Map of Diverse Immune Phenotypes in the Breast Tumor Microenvironment. Cell. 2018;174(5):1293–308 e36.

104. Zheng C, Zheng L, Yoo JK, Guo H, Zhang Y, Guo X, et al. Landscape of Infiltrating T Cells in Liver Cancer Revealed by Single-Cell Sequencing. Cell. 2017;169(7):1342–56 e16.

105. Tirosh I, Izar B, Prakadan SM, Wadsworth MH, 2nd, Treacy D, Trombetta JJ, et al. Dissecting the multicellular ecosystem of metastatic melanoma by single-cell RNA-seq. Science. 2016;352(6282):189–96.

106. Guo X, Zhang Y, Zheng L, Zheng C, Song J, Zhang Q, et al. Global characterization of T cells in non-small-cell lung cancer by single-cell sequencing. Nat Med. 2018;24(7):978–85.

107. Sathe A, Ayala C, Bai X, Grimes SM, Lee B, Kin C, et al. GITR and TIGIT immunotherapy provokes divergent multicellular responses in the tumor microenvironment of gastrointestinal cancers. Genome Med. 2023;15(1):100.

108. Clement K, Rees H, Canver MC, Gehrke JM, Farouni R, Hsu JY, et al. CRISPResso2 provides accurate and rapid genome editing sequence analysis. Nat Biotechnol. 2019;37(3):224–6.

109. Ayala C, Sathe A, Bai X, Grimes SM, Shen J, Poultsides GA, et al. Distinct gene signatures define the epithelial cell features of mucinous appendiceal neoplasms and pseudomyxoma metastases. Front Genet. 2025;16:1536982.

