## supplementary figures for "Colorectal cancer relies on an immunosuppressive cellular topography and genomic adaptations for establishing brain metastases"

**INSTITUTIONS:**

[1] Division of Oncology, Department of Medicine, Stanford University School of Medicine, Stanford, CA, USA

[2] Quantitative Sciences Unit, Department of Medicine, Stanford University School of Medicine, Stanford, CA, USA

[3]Department of Neurosurgery, Stanford University School of Medicine, Stanford, CA, USA

[4]Department of Neurosurgery, Johns Hopkins University School of Medicine, Baltimore, Maryland, USA

[5]Stanford Cancer Institute, Stanford University School of Medicine, Stanford, CA, USA

[6]Department of Pathology, Stanford University, Stanford, CA, USA

**CORRESPONDING AUTHOR**

Hanlee P. Ji

Mailing address: CCSR 2245, 269 Campus Drive, Stanford, CA-94305, USA



Conflict of interest statement: The authors declare no potential conflicts of interest.

#### SUPPLEMENTARY FIGURE LEGENDS

**Supplementary Figure 1:** (a) UMAP representation of Xenium data colored by patient. (b) Proportion of cell types across respective samples. (c-f) Scaled average expression of marker genes in in respective cell lineages.

**Supplementary Figure 2:** (a-b) UMAP representation of scRNA-seq reference colored by (a) datasets or (b) cell types. (c) Scaled average expression of lineage marker genes in respective cell types.

**Supplementary Figure 3:** (a) Proportion of cell types across respective samples. (b) Spearman correlation between Visium and Xenium counts for shared genes in respective cell lineages. (c) Representative spatial plots for selected gene expression in respective patients together with paired H&E.

**Supplementary Figure 4**: (a) Average scaled expression of respective genes in pure tumor spots from CRC-BMet or paired primary CRC from the same patient. (b) IHC staining for respective antibody or isotype control. (c) Volcano plot highlighting upregulated genes in pure tumor spots from CRC-BMet compared to CRC bladder metastasis from P2.

**Supplementary Figure 5:** (a) Average scaled GSVA score for respective hallmark pathway across tumor spots from all CRC-BMets.

**Supplementary Figure 6:** (a-d) Inferred CNV profiles from pure tumor spots from paired CRC-Bmets and primary CRC from respective patients, with suffix indicating subclone, together with CNV profiles from WGS.

**Supplementary Figure 7:** (a-b) Comparison of (a) CIN70 signature or (b) inferred FGA in pure tumor spots from CRC-BMet or CRC bladder metastasis from P2 with t-test *p*. (c) Inferred CNV profiles from paired bladder and CRC-BMet from P2, with suffix indicating subclone. (d) Average inferred CNV per chromosome arm pathway across tumor spots from all CRC-BMets.

**Supplementary Figure 8:** (a) Proportion of cell neighborhoods across respective samples. (b) Spatial representation of cellular neighborhoods in respective samples. Zoomed inset for P44 depicts IHC for respective proteins performed in adjacent sections.

**Supplementary Figure 9:** (a) Average importance score of cell-type abundance in the prediction of abundances of other cell types within a spot (65 μm). (b) Distribution of average importance score of tumor epithelial cell abundance in the prediction of fibroblasts and macrophage abundance within a spot across samples (65 μm) (c) Average importance score of cell-type abundance in the prediction of abundances of other cell types within adjacent spots (250 μm).

**Supplementary Figure 10:** (a) Proportions of spatial domains across patients. (b) Representation of spatial domains from respective patients together with paired H&E. (c) Upregulated genes in intermediate state tumor domain.

**Supplementary Figure 11:** (a-b) Upregulated pathways in respective spatial domain.

(c) Proportions of respective cell types between spatial domains together with Wilcoxon test *p*.

(d) Upregulated pathways in respective spatial domain.

**Supplementary Figure 12:** (a-c) Upregulated pathways in respective spatial domain. (d) Expression of respective genes in spatial domains. Size of the dot is proportional to the percentage of spots expressing the gene. (e) Scaled GSVA pathway activity across spatial domains.

**Supplementary Figure 13:** (a) Comparison of EMT signature in pure tumor spots from post-radiation and pre-radiation samples with t-test *p*. (b) IHC staining for respective antibody or isotype control. (c) Average scaled expression of respective genes in pure tumor spots from pre and post radiation samples from the same patient. (d-g) Inferred CNV profiles from pure tumor spots from paired pre or post radiation samples from respective patients, with suffix indicating subclone. (h) Comparison of proportions between paired pre- and post-radiation samples from the same patient. All comparisons have adjusted $p > 0.05$ using a one-sided Wilcoxon rank-sum test.

**Supplementary Figure 14:** (a-b) SPP1_KO1 induced deletion in comparison to ctrl gRNA with location and efficiency of editing. (c) Spearman correlation between scRNA-seq expression from co-cultures and Visium from CRC-BMet patient samples for each respective lineage. (d) Violin plots depicting expression of differentially expressed genes in tumor cells across co-culture conditions. (e) Violin plots depicting expression of differentially expressed genes in fibroblasts across co-culture conditions.

#### Supplementary figure 1

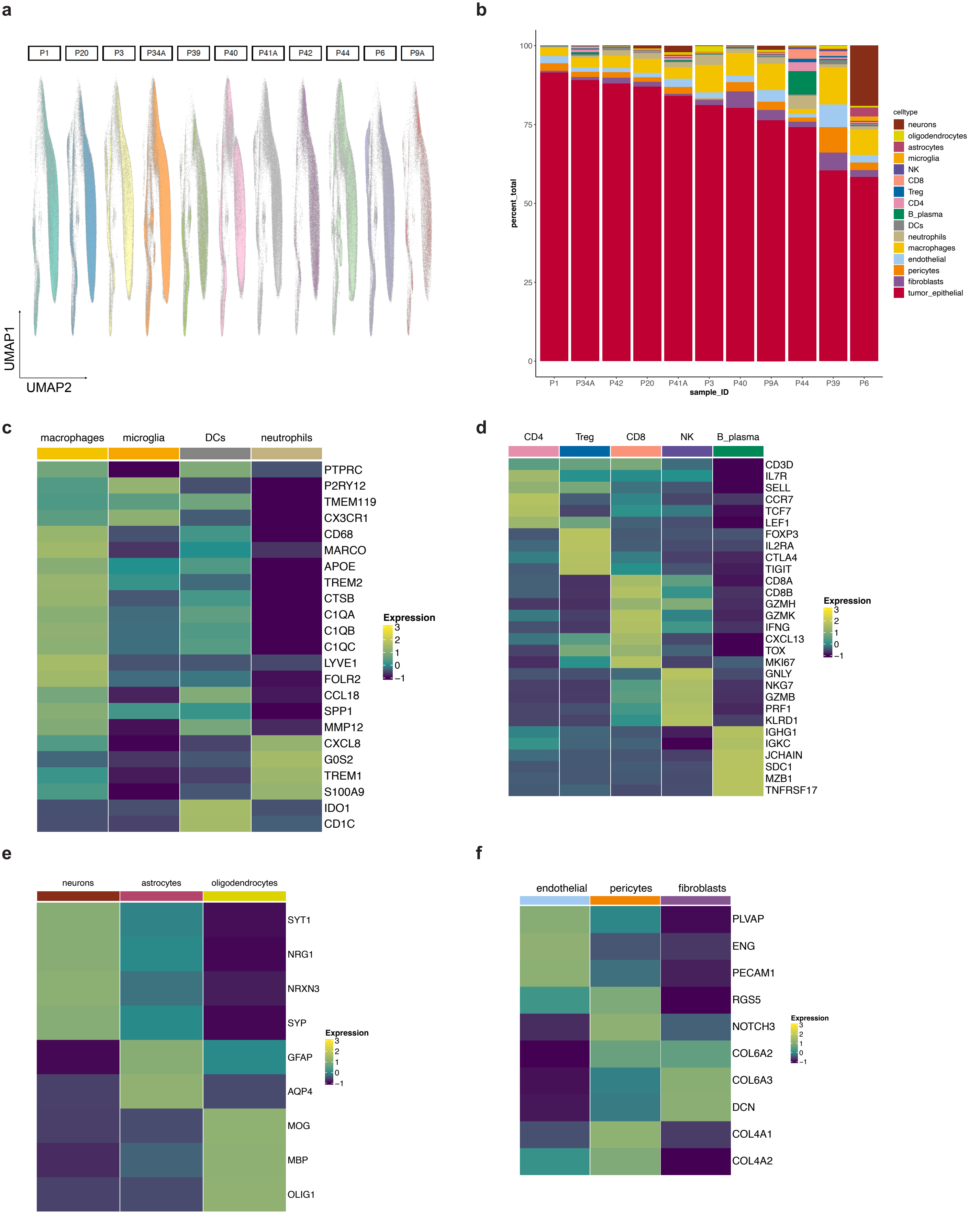

#### Supplementary figure 2

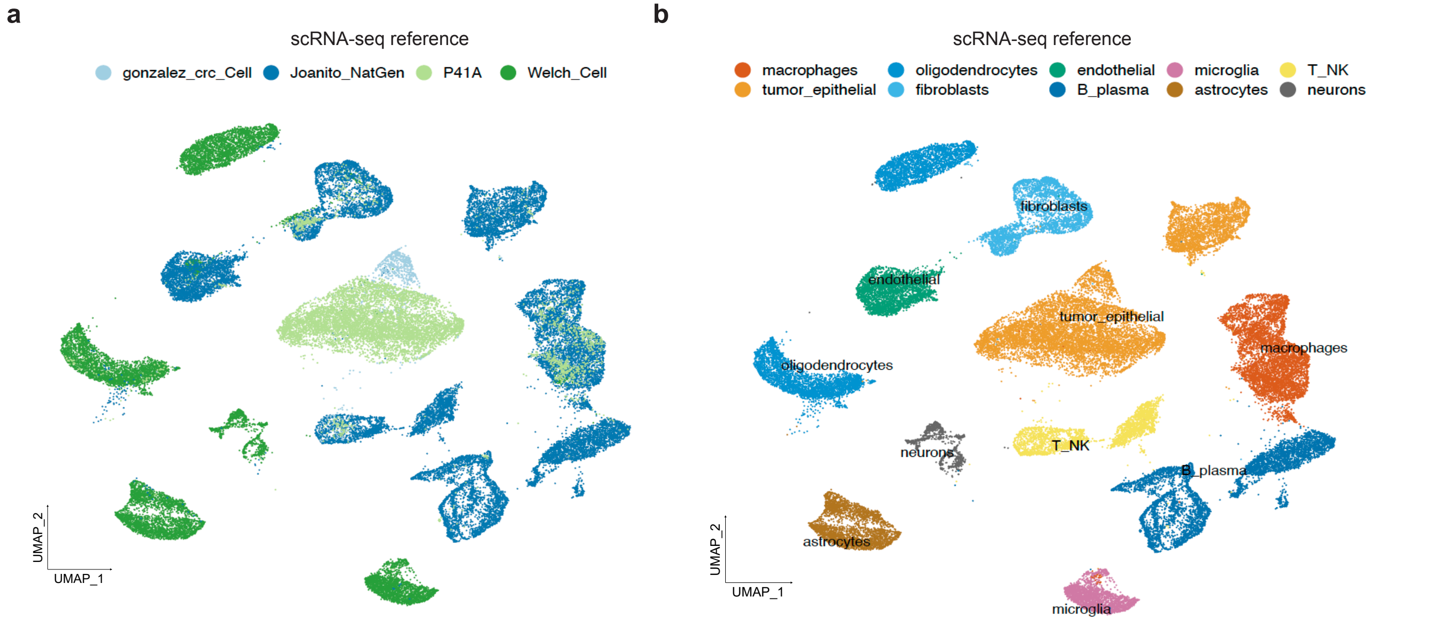


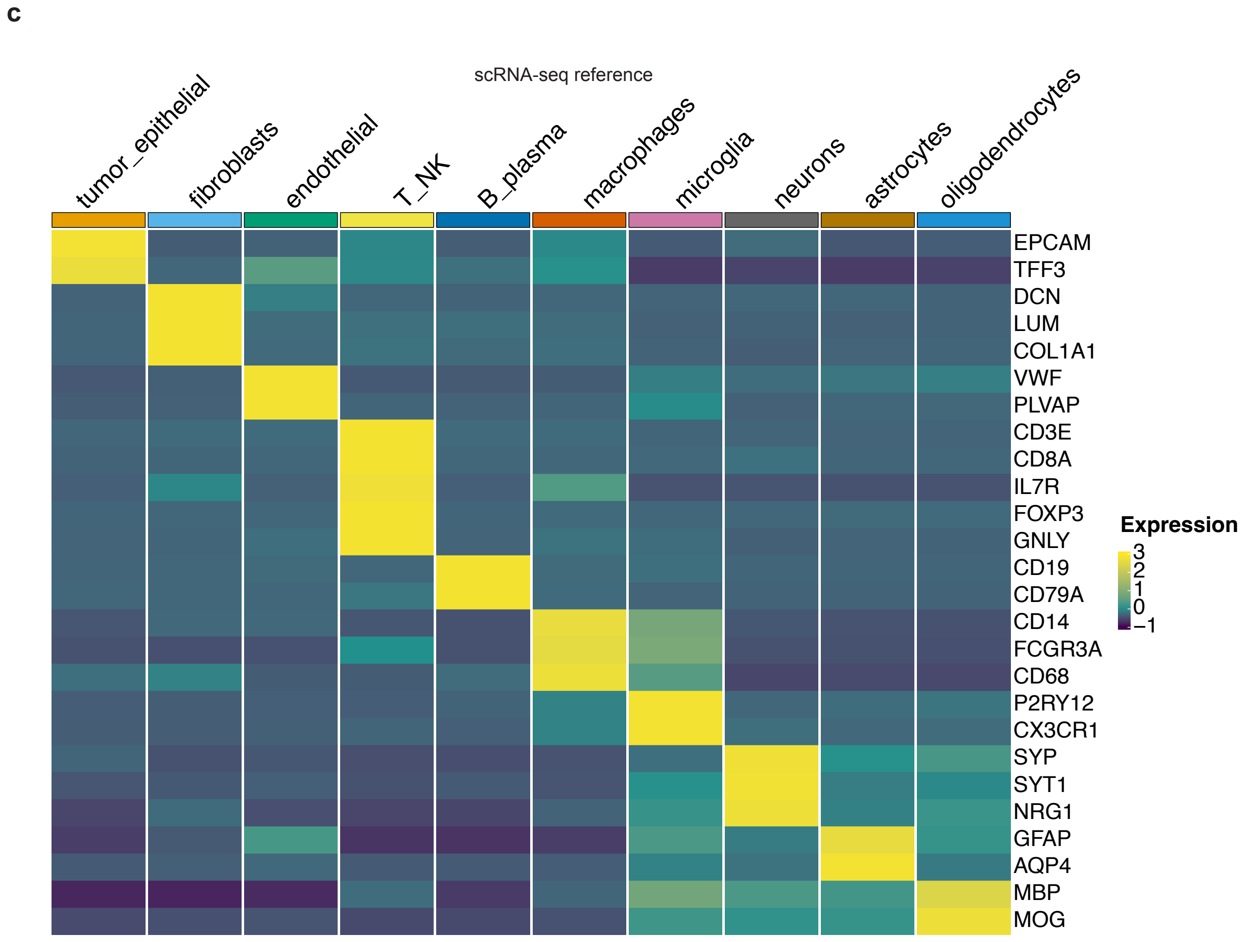

#### Supplementary figure 3

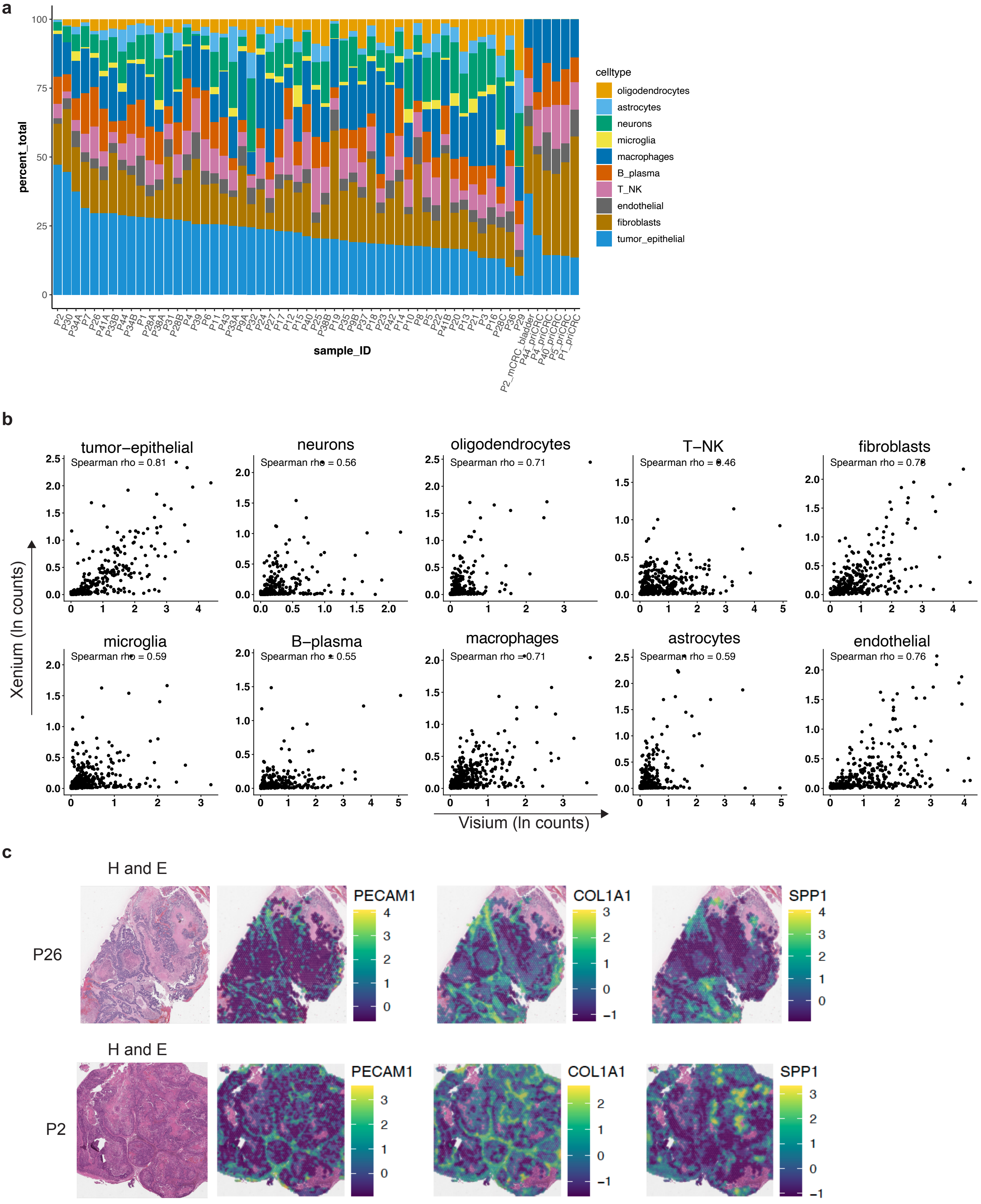

#### Supplementary figure 4

a

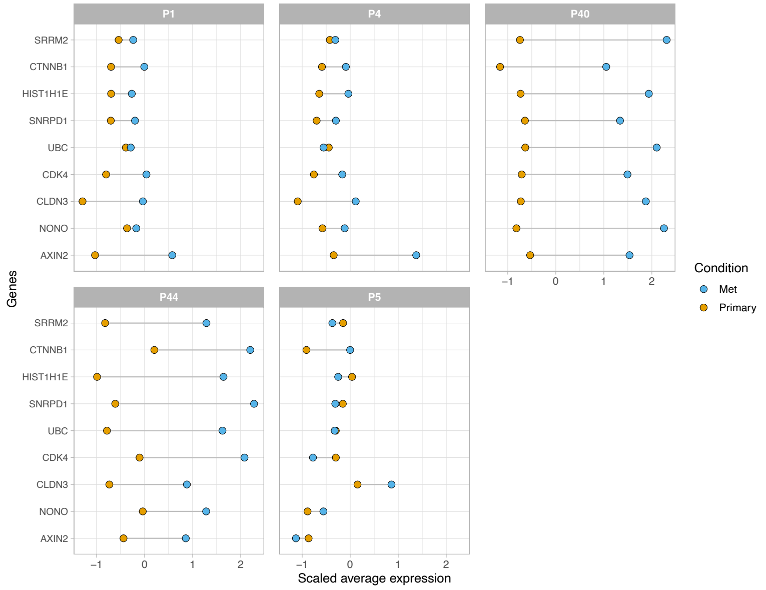


c

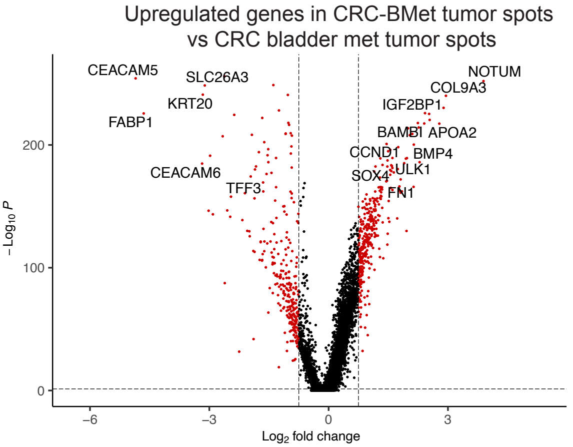


b

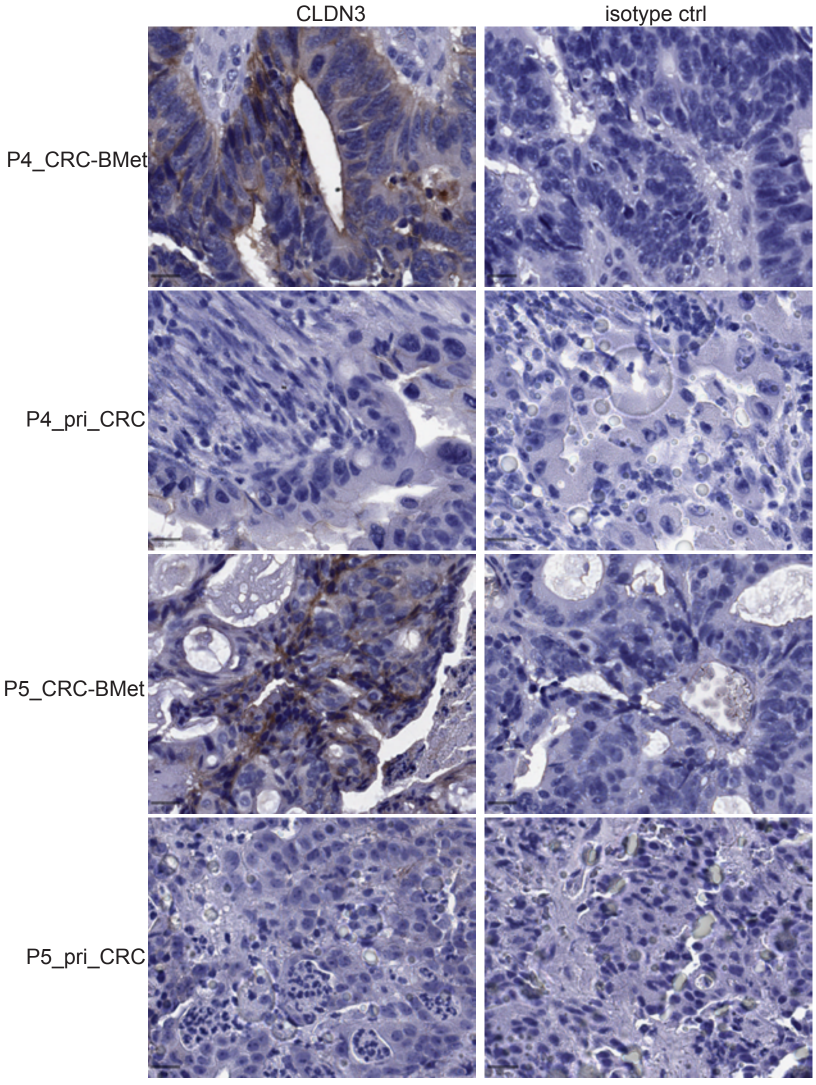

**Supplementary figure 5**

**a**

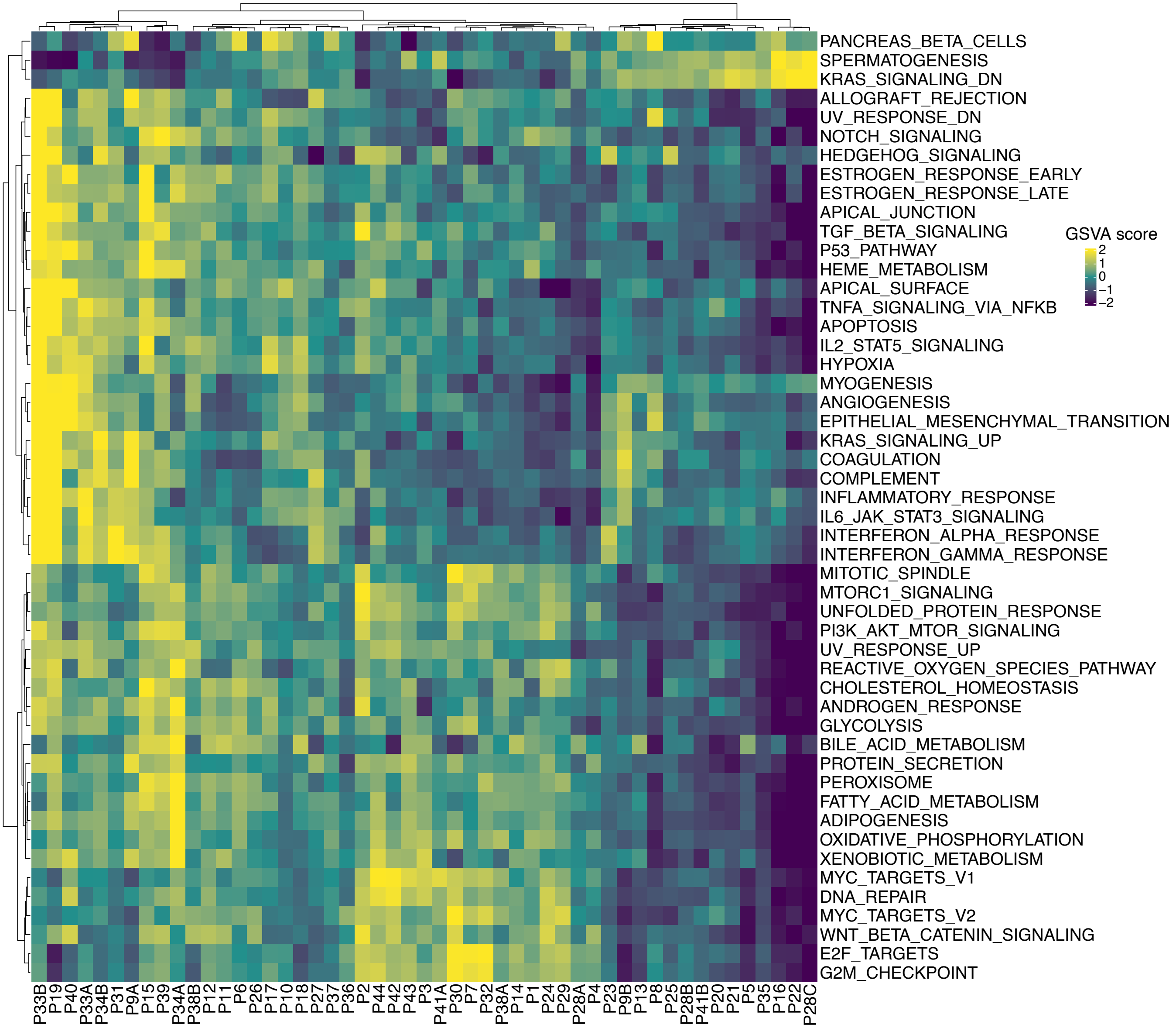

### Supplementary figure 6

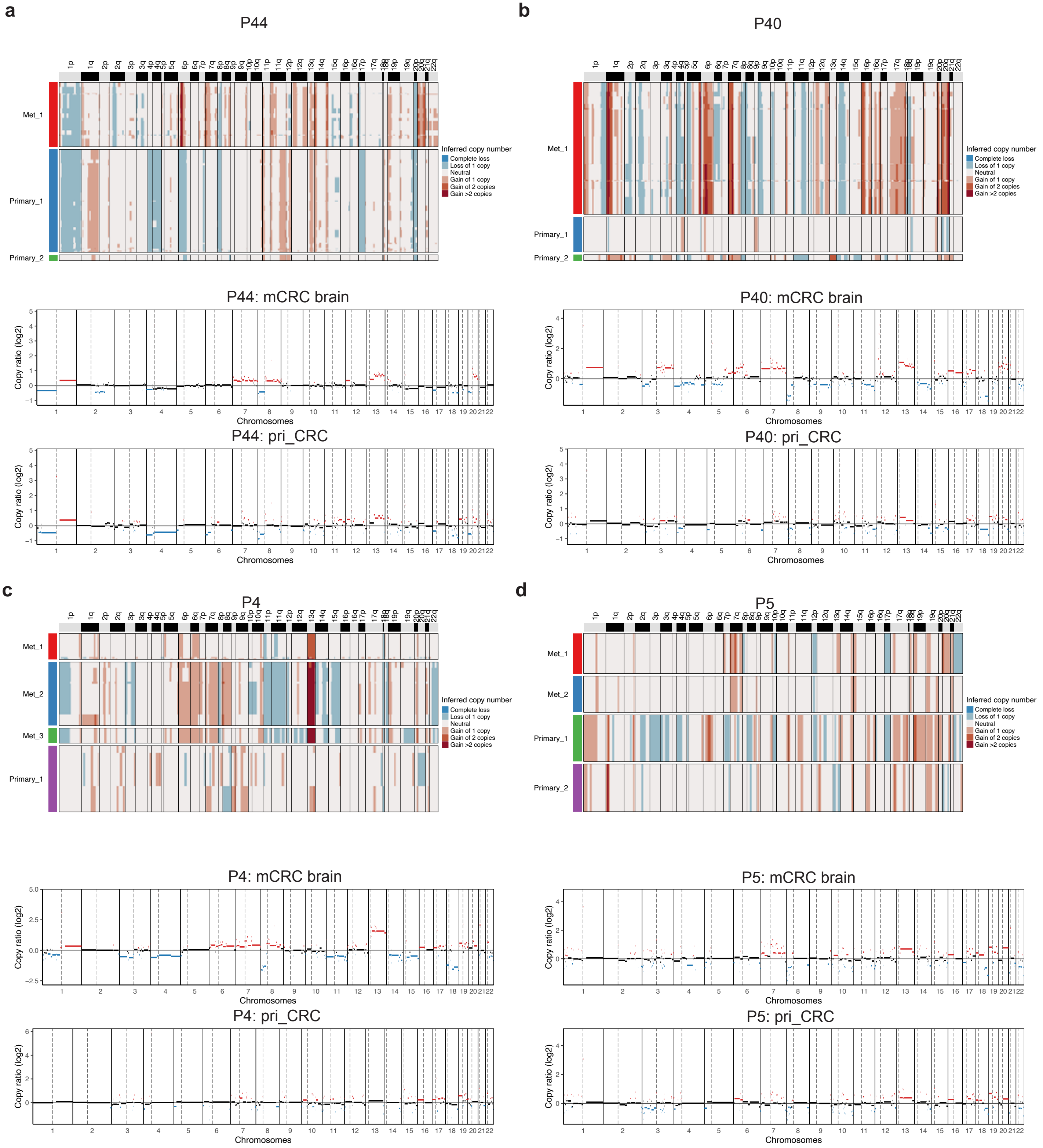

#### Supplementary figure 7

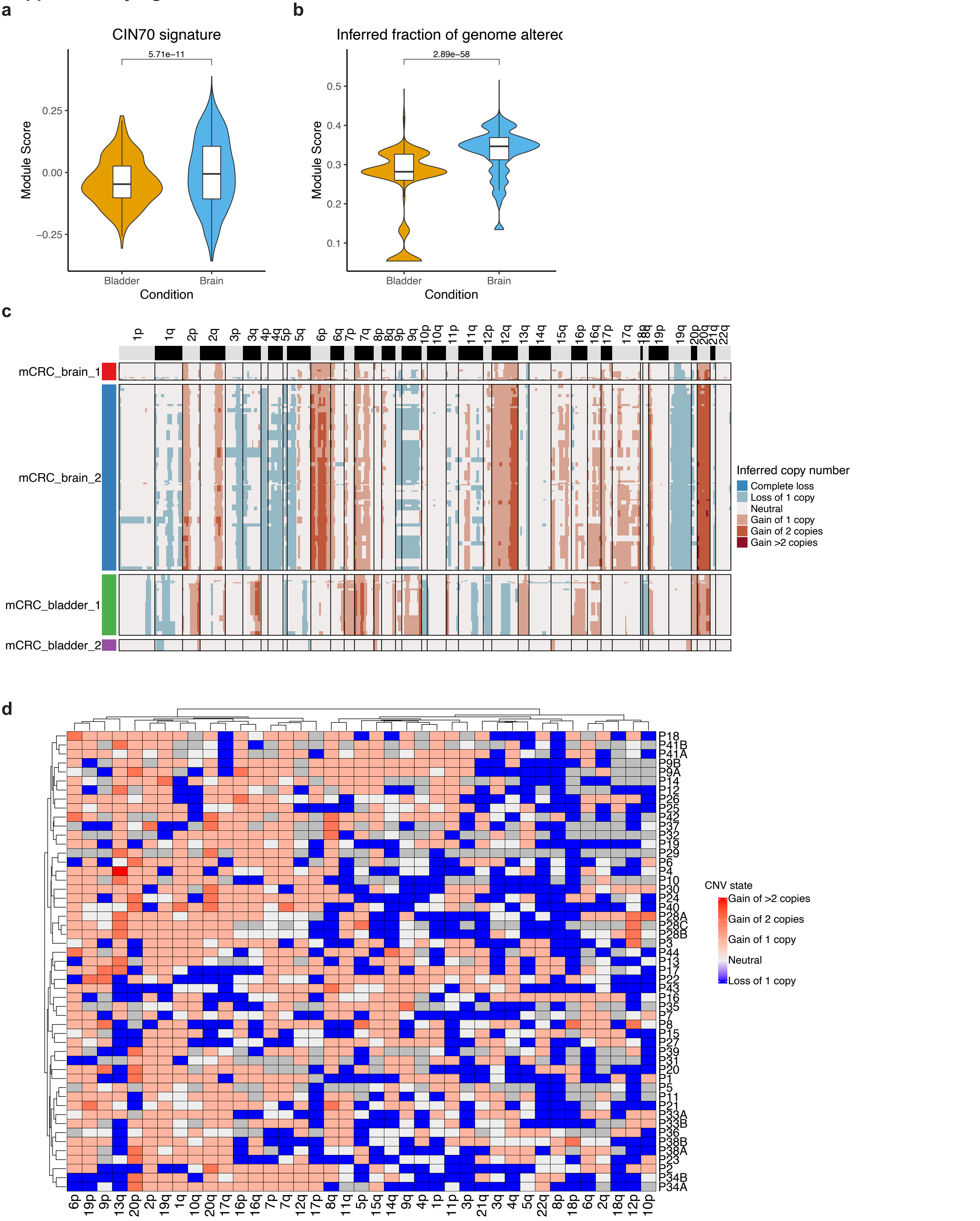

### Supplementary figure 8

**a**

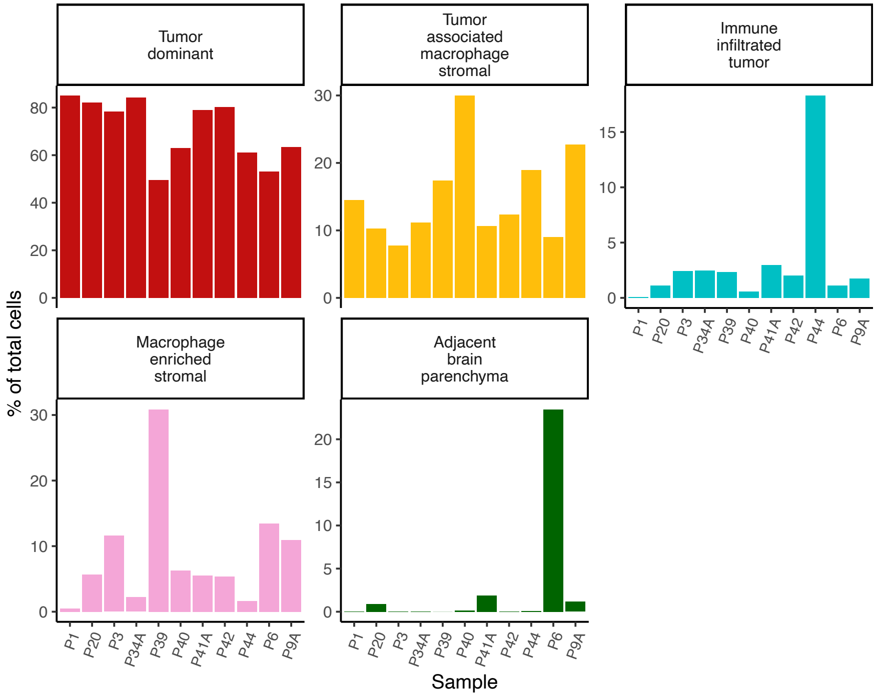


**b**

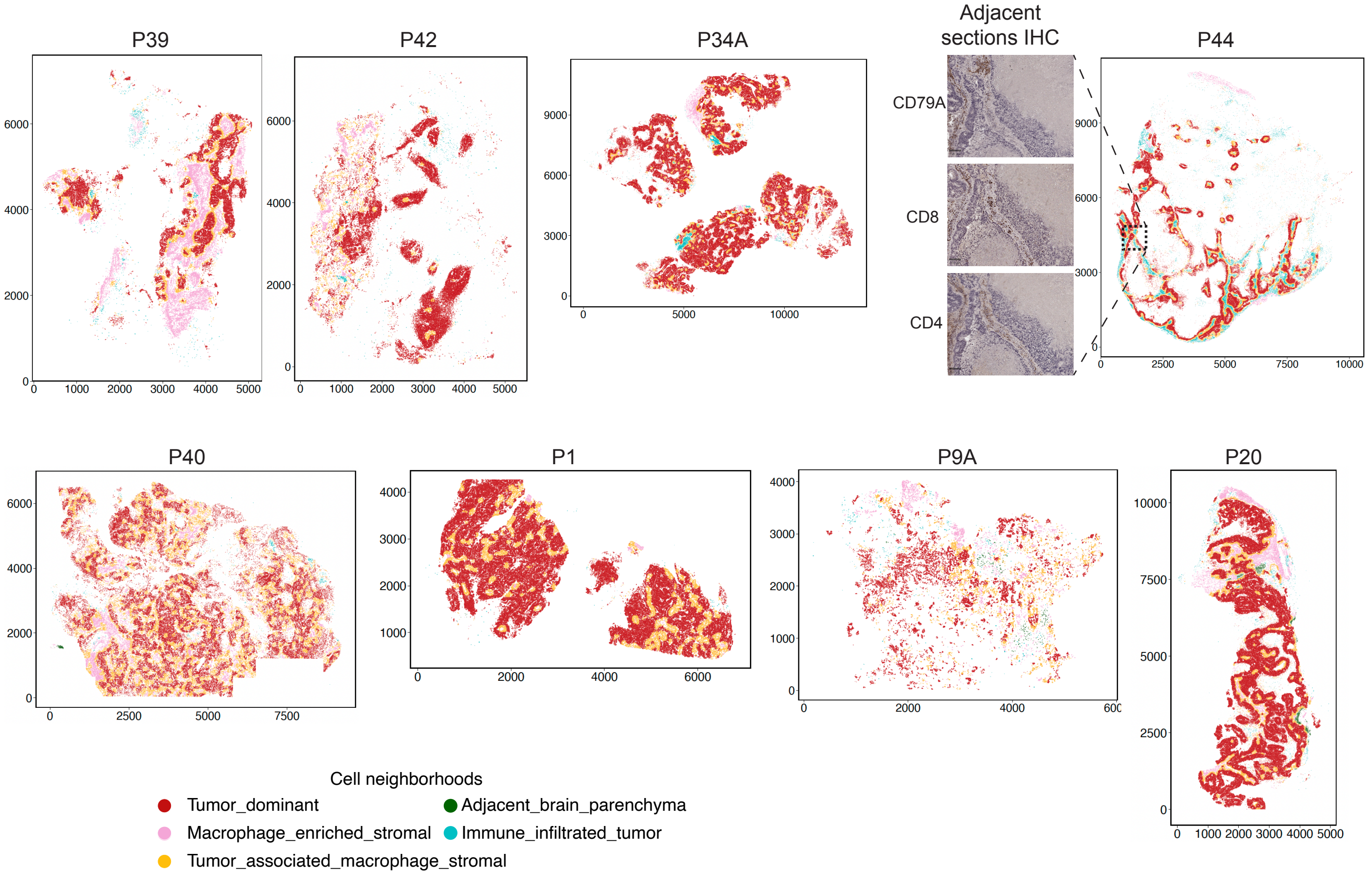

### Supplementary figure 9

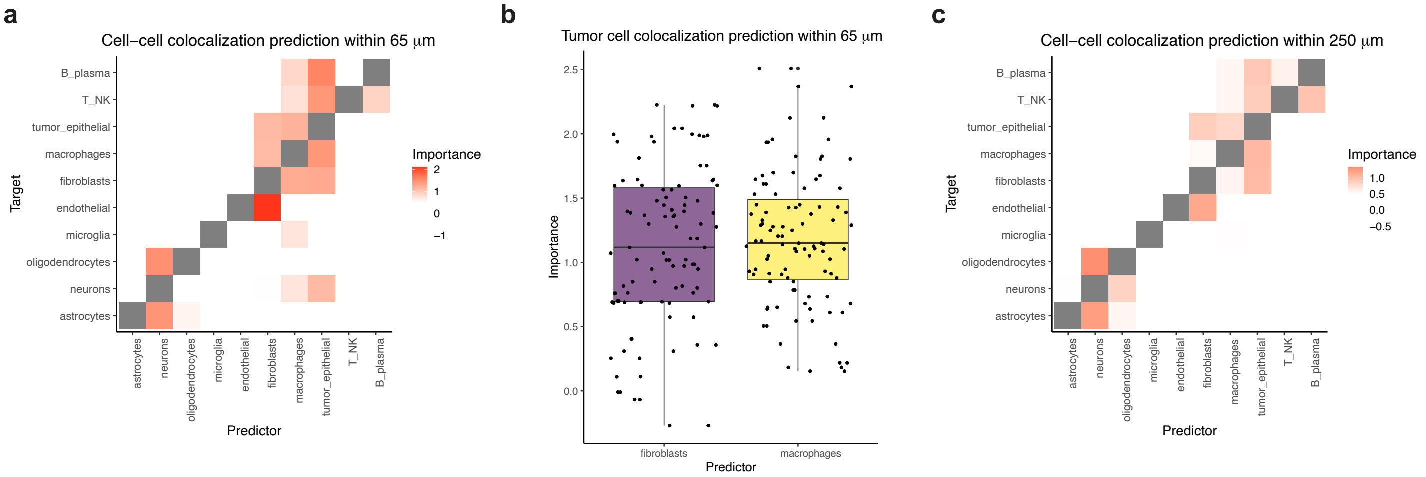

### Supplementary figure 10

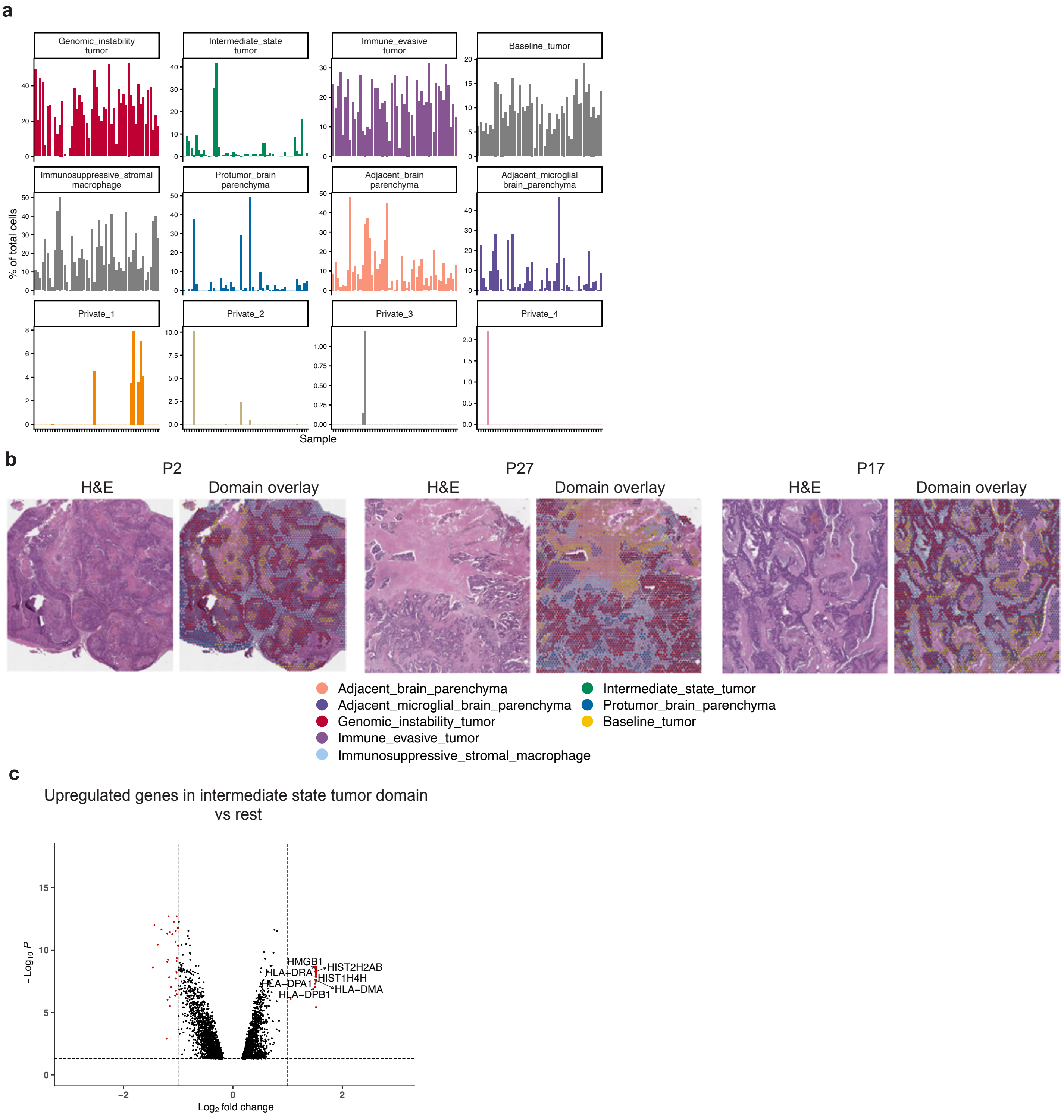

### Supplementary figure 11

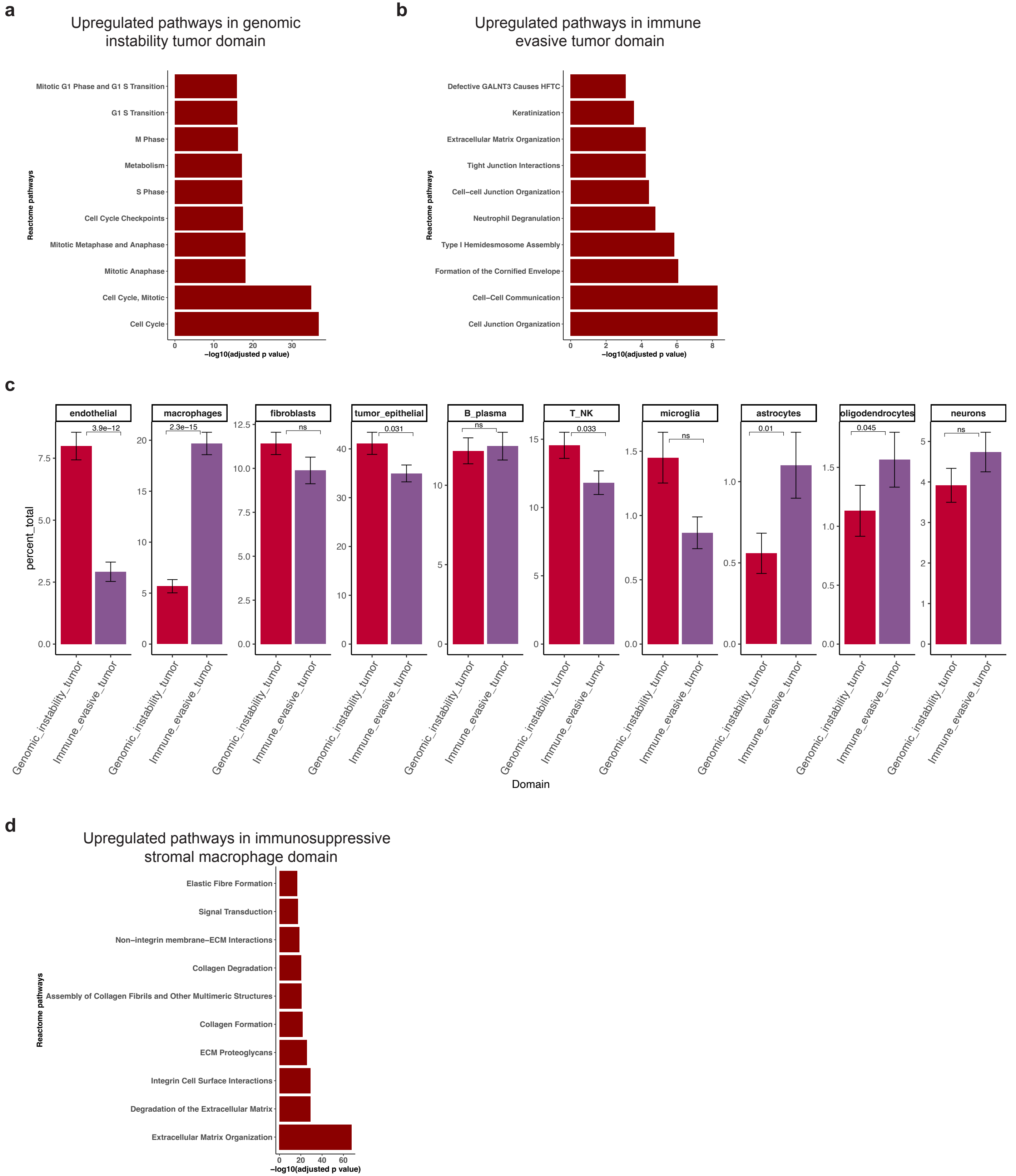

#### Supplementary figure 12

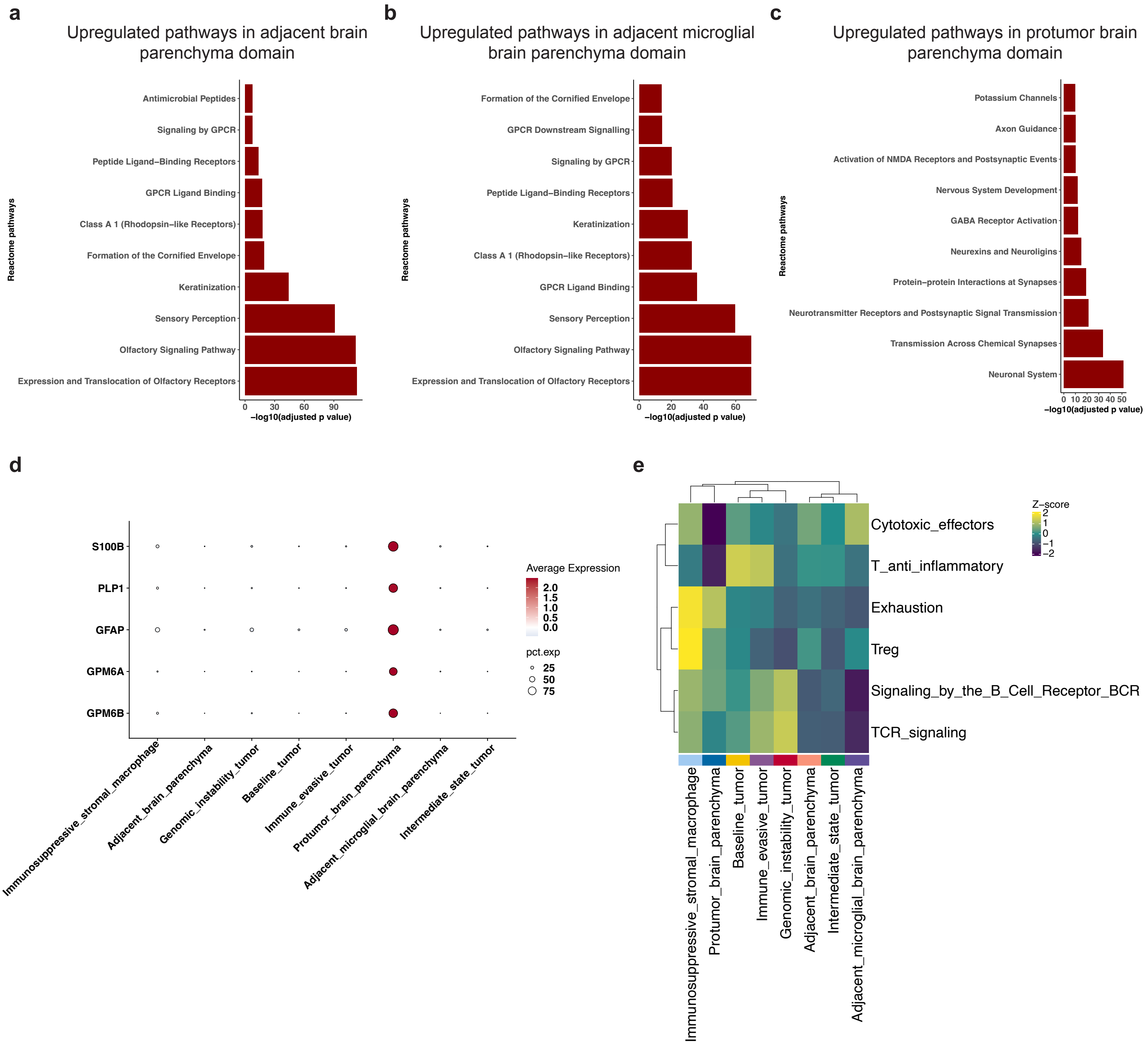

### Supplementary figure 13

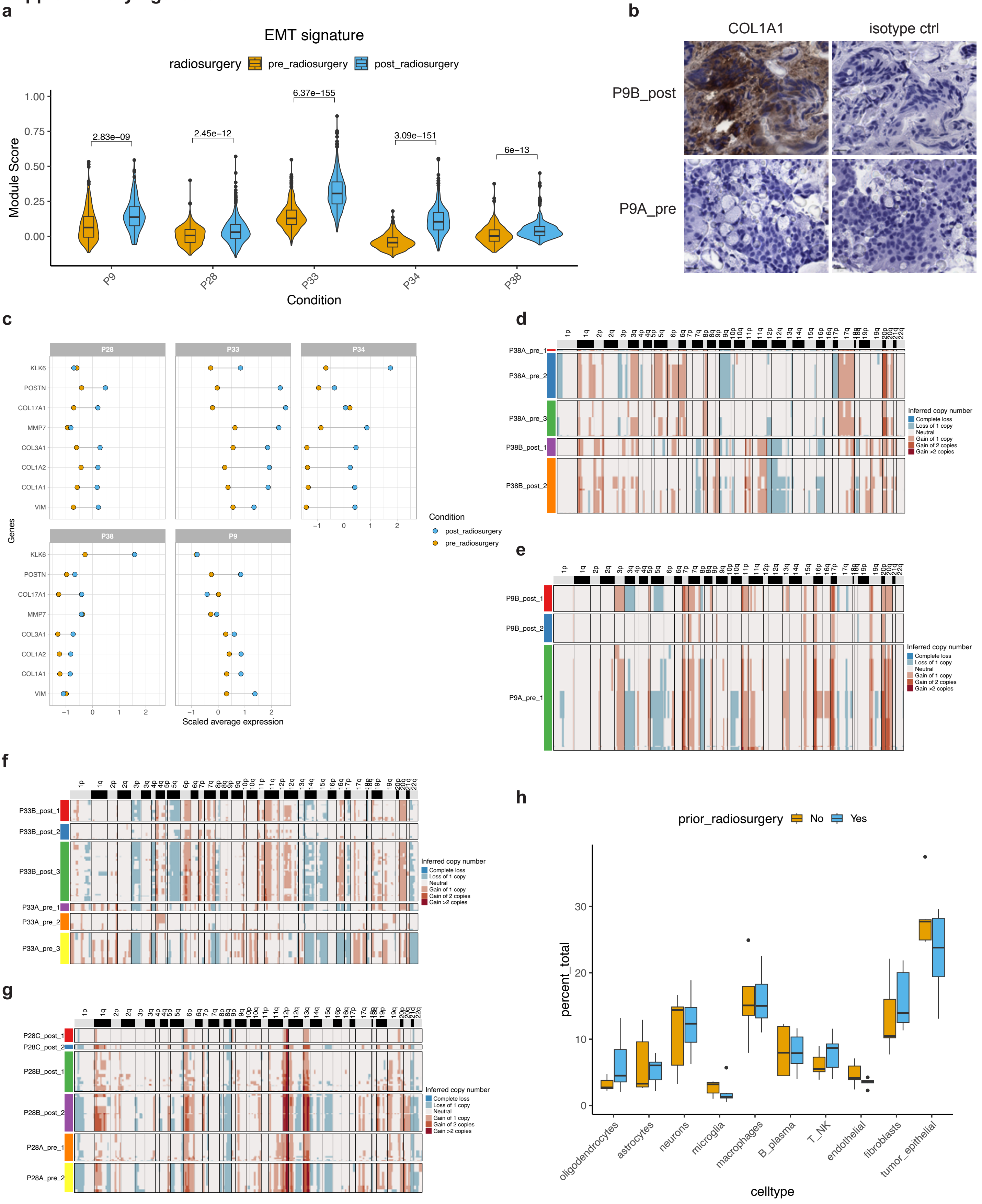

#### Supplementary figure 14

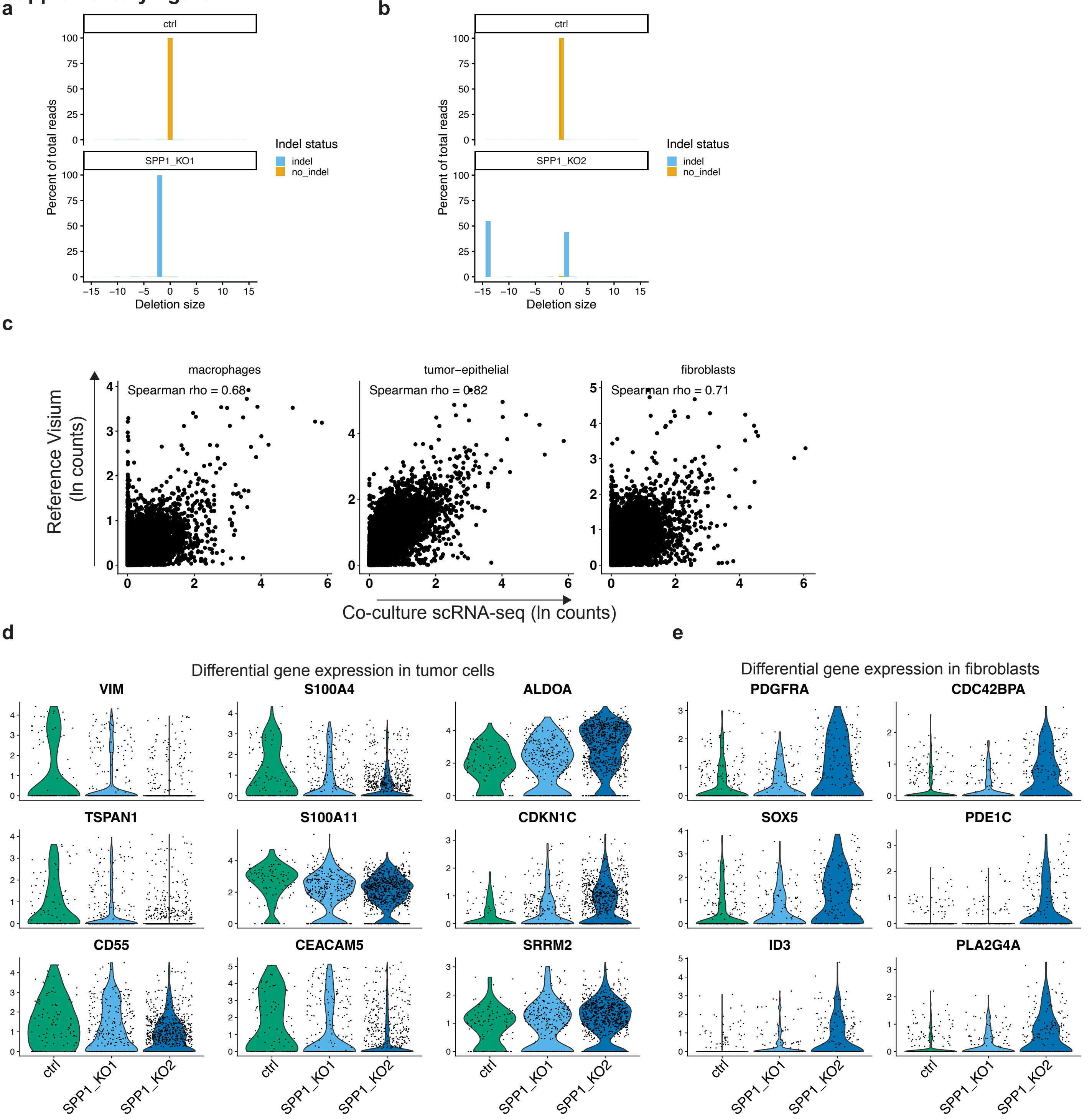
